# Genomic and Spectral Visual Adaptation in Southern Leopard Frogs during the Ontogenetic Transition from Aquatic to Terrestrial Light Environments

**DOI:** 10.1101/2021.02.19.432049

**Authors:** Ryan K Schott, Rayna C Bell, Ellis R Loew, Kate N Thomas, David J Gower, Jeffrey W Streicher, Matthew K Fujita

## Abstract

Many animals have complex life cycles where larval and adult forms have distinct ecologies and habitats that impose different demands on their sensory systems. While the adaptive decoupling hypothesis predicts reduced genetic correlations between life stages, how sensory systems adapt across life stages at the molecular level is not well understood. Frogs are a compelling system to study this question in because most species rely on vision as both aquatic tadpoles and terrestrial adults, but these habitats present vastly different light environments. Here we used whole eye transcriptome sequencing to investigate differential expression between aquatic tadpoles and terrestrial juveniles of the southern leopard frog (*Lithobates sphenocephalus*). Because visual physiology changes with light levels, we also tested how constant light or dark exposure affected gene expression. We found 42% of genes were differentially expressed in the eyes of tadpoles versus juveniles, versus 5% for light/dark exposure. Analyses targeting a curated set of visual genes revealed significant differential expression between life stages in genes that control aspects of visual function and development, including spectral sensitivity and lens composition. Light/dark exposure had a significant effect on a smaller set of visual genes. Finally, microspectrophotometry of photoreceptors confirmed shifts in spectral sensitivity predicted by the expression results, consistent with adaptation to distinct light environments. Overall, we identified extensive expression-level differences in the eyes of tadpole and juvenile frogs related to observed morphological and physiological changes through metamorphosis, and corresponding adaptive shifts to optimize vision in aquatic versus terrestrial environments.

## Introduction

Many animals occupy different ecological niches throughout their life cycles, which can result in distinct selective pressures at different developmental stages (Werner 1988; Ebenman 1992). Animals with simple life cycles may be phenotypically constrained due to genetic correlations between juvenile and adult morphologies, whereas complex life cycles in which animals develop through two or more distinct phenotypic phases may disrupt these constraints through adaptive decoupling (Ebenman 1992). This hypothesis predicts that each life stage can adapt more freely to its particular ecological niche (Ebenman 1992; Moran 1994). Adaptive decoupling has been documented in different taxa, including marine invertebrates, insects, and amphibians, and across many types of traits, including morphology, colouration, and social behaviour (Moran 1994; Anderson, et al. 2015; Sherratt, et al. 2017; Medina, et al. 2020). The degree of decoupling can vary considerably, and in many cases larval trait characteristics heavily influence how those traits manifest in adults (Azumi, et al. 2007; Crean, et al. 2011; Marshall and Morgan 2011; Aguirre, et al. 2014; Bonett and Blair 2017; Phung, et al. 2020). Investigating significant changes in gene expression across life stages can be an important step towards understanding the genetic basis of adaptive decoupling (Collet and Fellous 2019). Our study provides important insight into the molecular basis of adaptive decoupling in the visual system by quantifying differences in gene expression profiles across life stages in frogs.

The adaptive decoupling hypothesis has been studied in anuran amphibians (frogs and toads) due to their complex life cycles that typically include distinct larval (tadpole) and adult stages linked by metamorphosis. Although both tadpole and adult ecologies vary extensively across the frog tree of life, most species have aquatic and herbivorous tadpoles, while adults are generally more terrestrial and carnivorous (McDiarmid and Altig 1999). A number of studies have shown that morphological diversity of tadpoles and adult frogs is decoupled (Roelants, et al. 2011; Sherratt, et al. 2017; Wollenberg Valero, et al. 2017; Fabrezi, et al. 2019) and that genetic correlations between tadpole and adult traits are low in many, but not all, cases (Blouin 1992; Phillips 1998; Watkins 2001; Goedert and Calsbeek 2019). However, several aspects of frog biology do tend to be coupled across life stages to varying degrees including behavior (Wilson and Krause 2012), size (Dehling and Sinsch 2019; Phung, et al. 2020), and developmental plasticity (Trokovic, et al. 2011). In general, the degree of coupling/decoupling across anuran life stages is trait, and possibly taxon, dependent.

The visual system of frogs is a particularly compelling trait in the context of adaptive decoupling because most species use vision to sense their environments as both tadpoles and adults, but differences in morphology, ecology, and behavior between life stages likely place different selective pressures on the visual system across ontogeny. Correspondingly, intraspecific eye-body size allometry across ontogeny varies widely across anurans with a shift at metamorphosis in several species, suggesting that eye growth is partially decoupled and is shaped by both tadpole and adult visual requirements (Shrimpton et al. *in press*). Eye position also shifts in many anurans from a lateral position in tadpoles to a more frontal position in adults, resulting in binocular overlap (Hoskins 1990). Other morphological and physiological changes to the visual system that may also occur between larval and adult life stages in anurans include the loss or development of accessory structures (e.g., umbracula, elygia, eyelids, nictitating membranes), changes in photoreceptor and ganglion cell morphology, and shifts in synaptic connections. As with most changes associated with metamorphosis, changes to the visual system are broadly controlled by thyroid hormone (Hoskins 1990; Mann and Holt 2001). The specific molecular basis of these broad morphological and physiological changes to the visual system, however, are poorly understood, especially in species that transition from aquatic tadpoles to terrestrial adults.

Aquatic and terrestrial environments differ in both the intensity and spectral composition of available light, and thus, the visual systems of anurans with aquatic larvae and terrestrial adults can encounter vastly different light environments. Water preferentially absorbs and scatters the shorter (ultraviolet–violet) and longer (yellow–red) wavelengths of the light spectrum resulting in a narrowing of the spectrum and overall reduced light availability. In clear water this results in a depth-dependent blue-shift in available light (Lythgoe 1979). Freshwater environments often have dissolved organic and particulate matter that absorbs shorter (violet and blue) wavelengths, resulting in a red-shifted light environment, and these particles may also further reduce the penetration of light with depth (Levine and MacNichol 1982; Costa, et al. 2013). Many vertebrate animals that inhabit turbid, red-shifted aquatic environments use visual pigments that have red-shifted sensitivity (relative to species in marine or terrestrial environments) that presumably match the available light more closely (Wald 1939; Bridges 1972; Toyama, et al. 2008). This shift can be accomplished through the use of a light-sensitive chromophore derived from vitamin A2 (3,4-didehydroretinal) in contrast to “typical” vertebrate visual pigments that contain a vitamin A1 derived chromophore (retinal). Consequently, some frog species that transition from an aquatic tadpole to a terrestrial adult have a corresponding shift in chromophore usage from mainly A2 to predominantly, or exclusively, A1 (Bridges 1972). By contrast, African clawed frogs, *Xenopus laevis*, are fully aquatic throughout their lifecycle and exclusively use the A2 chromophore (Bridges, et al. 1977). The conversion from the A1 to A2 chromophore is mediated by a cytochrome enzyme encoded by *CYP27C1* (Enright et al. 2015), and thus differential expression of this gene in aquatic versus terrestrial life stages likely plays an important role in optimizing visual sensitivity in these distinct light environments.

Differential visual opsin usage and expression is another mechanism of adaptation to the changing light conditions that tadpoles and adult frogs experience across life stages. Visual opsins are the protein component of visual pigments, and changes to the protein sequence can affect spectral sensitivity. Vertebrates ancestrally have five classes of visual opsins (LWS, RH1, RH2, SWS1, SWS2) that are sensitive to different portions of the visual spectrum, although gene loss and duplication are common in some lineages (Wayne, et al. 2012). Frogs, for instance, have lost RH2, whereas teleost fishes often have additional, duplicated copies of some opsin classes (Carleton, et al. 2020). Differential visual opsin usage and expression across life stages is employed by many teleosts, particularly when larval habitat or foraging differs from that of adults (Carleton, et al. 2020). For example, Midas cichlids express *SWS1* only early in ontogeny and express *SWS2* only, or primarily, late in ontogeny. Expression of other opsin genes also varies significantly over ontogeny (Härer, et al. 2017). Whether anurans also use this strategy has yet to be investigated.

Aquatic and terrestrial environments also have different optical properties that apply divergent selective pressures on the lens. In water, the cornea (the outer casing of the eye) has little to no focusing power due to the similar refractive indices of water and the fluid within the eye (aqueous humor; Land and Nilsson (2012)). As a result, the lens alone is responsible for focusing images on the retina in aquatic settings. By contrast, in air the cornea has substantial focusing power that varies based on its curvature, and thus a high powered lens suitable for an aquatic environment would result in over-focusing (Land and Nilsson 2012). In anurans with terrestrial adult life stages, the lens becomes flatter during metamorphosis, reducing its power (Sivak and Warburg 1983; Mathis, et al. 1988; Hoskins 1990), whereas lens shape changes little in frogs that remain aquatic as adults (Chung, et al. 1975). The protein composition (crystallins) of the anuran lens may also change during ontogeny, although this appears to occur as lens diameter increases rather than specifically at metamorphosis (Smith-Gill and Carver 1981). For instance, *Lithobates pipiens*, *L. catesbianus*, and *X. laevis* show a shift from predominantly gamma crystallins to alpha and beta crystallins as eye and lens diameter increase (Polansky and Bennett 1973; Doyle and Maclean 1978; Hoskins 1990). Thus, genes that regulate lens growth and composition may also be differentially expressed through ontogeny, and this has yet to be explored in anurans.

Finally, the plasticity of visual gene expression in larval and adult anurans, as well as other non-model vertebrates, is poorly understood. In particular, the effect of short-term light or dark exposure is one axis of variation that may be especially important to consider with respect to experimental design in vision research. Animals are often exposed to dark conditions for several hours (i.e., dark adapted) prior to sampling to aid in dissection of the retina and to ensure photoreceptors have not been bleached (activated). The isolated retina is then used for downstream applications such as RNA sequencing or microspectrophotometry (MSP). By contrast, studies that use whole eyes to assess gene expression (e.g., whole eye transcriptome sequencing), may not use dark adaptation prior to sampling and in general may have more variable sampling conditions, especially when individuals are sampled in the field. Consequently, variability in visual gene expression with light conditions could have important, unappreciated implications for comparative studies of visual evolution.

Here we used whole eye transcriptome sequencing of the southern leopard frog (*Lithobates sphenocephalus*) to test for differential expression between fully aquatic tadpoles and post-metamorphic, terrestrial juveniles. Specifically, we 1) make broad transcriptome-wide comparisons to test for potential adaptive decoupling in gene expression between tadpole and juvenile eyes, and plasticity in response to light exposure, 2) use a curated subset of visual genes to specifically test for differential expression between life stages and light treatments, and 3) use MSP to analyze spectral sensitivity of tadpole and adult photoreceptor cells to bolster the conclusions we draw from the results of our differential expression analyses. This study provides a first look at how molecular aspects of the visual system change in a vertebrate that transitions from an aquatic to a terrestrial light environment and how variable these changes are with respect to light conditions during sampling.

## Results

### Transcriptome Sequencing and Assembly

Sequencing resulted in an average of 33 million paired end reads per sample, which was reduced to 22 million on average after quality control (Supplementary Table S1). The reference transcriptome assembled *de novo* with reads from all 12 samples resulted in 684,947 Trinity transcripts with an N50 of 1065, a 98.9% overall realignment rate, and high completeness, with 88.6% of BUSCOs complete and a further 4.2% of BUSCOs fragmented (Schott et al. 2021). A PCA plot identified one of the 12 samples as a conspicuous outlier (Supplementary Fig. 1). Potential explanations for this outlier include a range of factors that are unrelated to the aims of our study including variation in age, size, or condition among our field-caught samples, as well as contamination and/or RNA degradation during dissection and processing of the tissue. Consequently, the outlier was removed and we used the remaining 11 samples to produce an updated reference transcriptome with 634,894 Trinity transcripts, an N50 of 1085, the same re-alignment rate as the initial transcriptome (98.9%), and a slightly lower BUSCO completeness with 87.4% BUSCOs complete and a further 4.9% of BUSCOs fragmented (Schott et al. 2021). The transcriptome was reduced to a ‘best set’ of transcripts (reduced transcript set; see Methods and Schott et al. 2021), which resulted in a reduction to 66,165 transcripts with an N50 of 2077 and a slight drop in BUSCO scores (87.1% complete and 4.6% fragment), but a substantial drop in the number of duplicated BUSCOs (50.1% to 6.9%) indicating a substantial reduction in the redundancy of the transcriptome.

### Transcriptome-wide Analyses Reveal Substantial Differential Expression between Life Stages

A PCA of rlog transformed counts showed strong separation of the samples based on life stage (tadpole vs juvenile), but not light vs dark exposure (Fig. 1). Differential expression between tadpoles and juveniles was detected in 11,046 out of 23,019 transcripts (42% significant with an adjusted p-value < 0.05) with nonzero total read counts (Fig. 2, Supplementary File 1). By contrast, only 122 transcripts (5% of total) were differentially expressed between light and dark exposure (Fig. 2, Supplementary File 1). Of the 11046 differentially expressed transcripts between tadpoles and juveniles, 9038 were annotated with a total of 6721 GO terms (Schott et al. 2021). Twenty-four GO terms were found to be enriched (p < 0.01) with the top three terms being translation, retinol metabolic process, and regulation of small GTPase mediated signal transduction (Supplementary Fig. 2, Supplementary File 2). While retinol metabolic process, retinoic acid metabolic process, and regulation of synapse structure or activity have implications for visual system development, most other significant terms are likely related to general physiological differences between tadpoles and adults (e.g., oxygen transport, carbohydrate metabolic process, cell morphogenesis). Three other terms related to the visual system development were significant at the 0.05 level (positive regulation of neural retina development, p = 0.026; photoreceptor cell morphogenesis, p = 0.044; retinal pigment epithelium development, p = 0.050). GO terms related directly to visual function were not significant, but these were annotated with relatively few terms. For example, only 71 transcripts were annotated as visual perception and 11 as phototransduction (Supplementary File 2). This is likely the result of relying on a *de novo* assembly and *X. laevis* for annotation, and so we focused on our curated set of visual genes for the remainder of the analyses.

**Figure 1.**
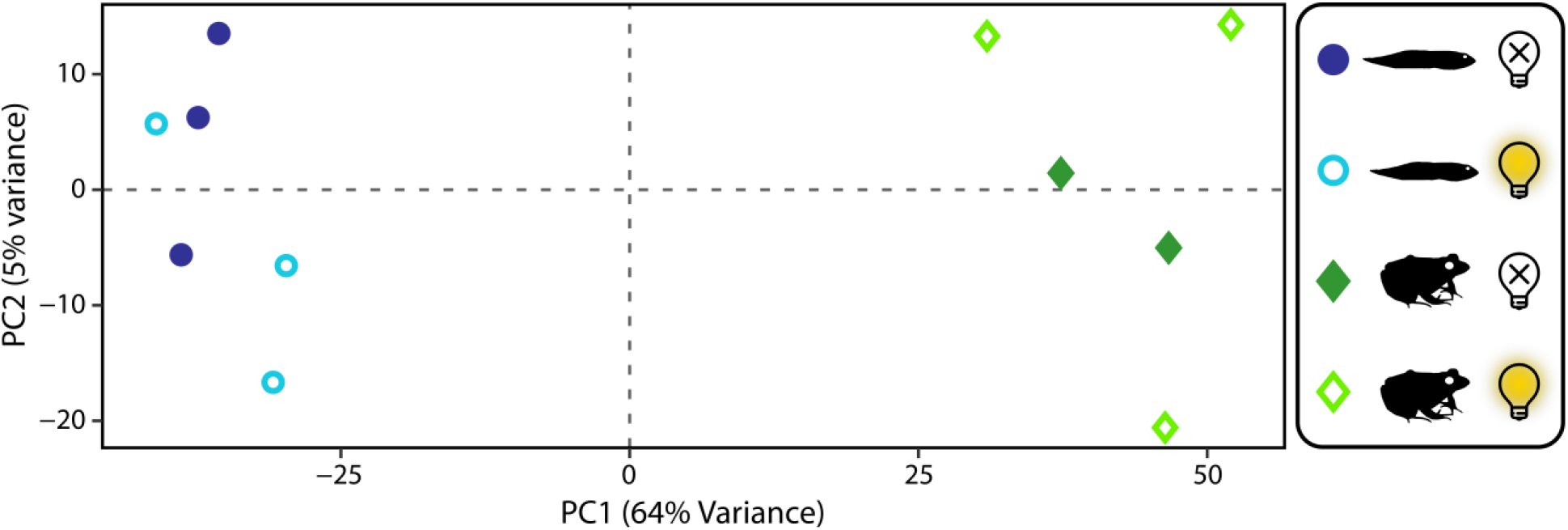
Principal components analysis plot of rlog transformed counts. The first principle component (PC1) accounts for 64% of the variance and clearly separates juveniles and tadpoles. Light and dark exposure are not clearly separated by PC2, which accounts for only 5% of the variance. One of the dark-exposed, juvenile samples was found to be an outlier and was removed prior to this analysis but is shown in Supplementary Figure 1.

**Figure 2.**
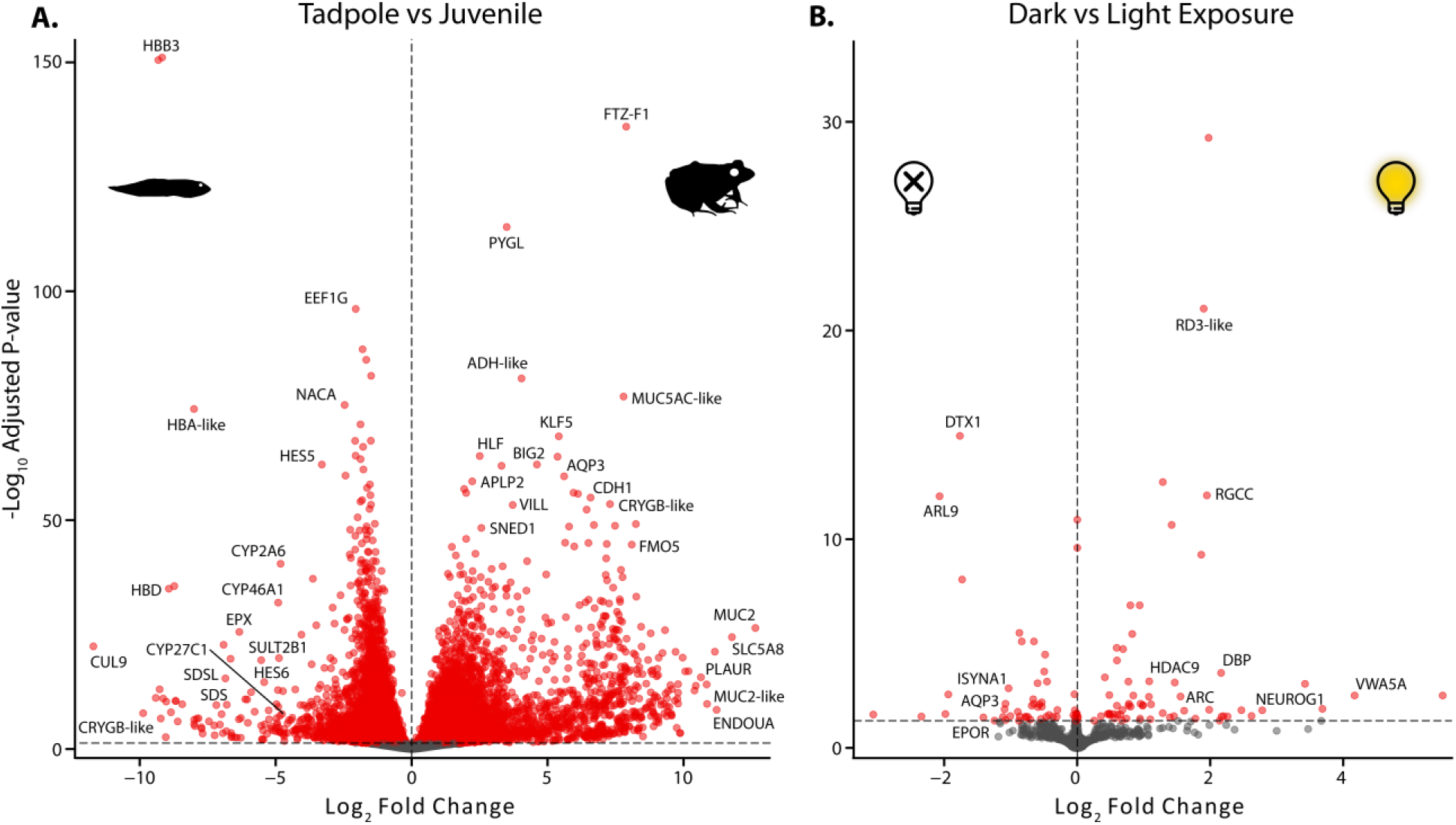
Volcano plots of transcriptome-wide differential expression between **A)** tadpoles and juveniles and **B)** dark and light exposure. Differential expression was estimated with DESeq2 using a multifactor design that accounted for the variation both in the life stages (tadpole vs juvenile) and treatments (dark vs light exposure). For visualization, log_2_ fold changes (LFC) were shrunk and an adjusted p-value of 0.05 was set as the significance cutoff.

### Visual Genes are Significantly Differentially Expressed between Life Stages and Light Treatments

To analyze variation specific to the visual system, we generated a second differential expression dataset using a curated set of 170 visual gene coding sequences that affect numerous aspects of visual function, including eye and retinal development, light detection, lens crystallins, and phototransduction (Supplementary File 3; Schott et al. 2021). A PCA of rlog transformed counts showed strong separation of the samples based on lifestage (tadpole vs juvenile), but not light vs dark exposure, similar to the transcriptome-wide results (Supplementary Fig. 3). We found 111 genes (69%) that were identified as being differentially expressed between life stages with an adjusted p-value < 0.05 (Fig. 3, Supplementary File 4). This percentage of significant differential expression is higher than that found transcriptome-wide and may be so, in part, because only functionally relevant visual genes were included in the curated dataset. Several types of visual genes had strong support for differential expression between tadpoles and juveniles, including chromophore usage, visual opsin, and lens crystallin genes that have clear consequences for vision in different light environments and are addressed in more detail below. In addition, tadpoles and juveniles differed in expression of visual and photoreceptor development genes, several non-visual opsins, and a number of phototransduction and visual cycle genes.

**Figure 3.**
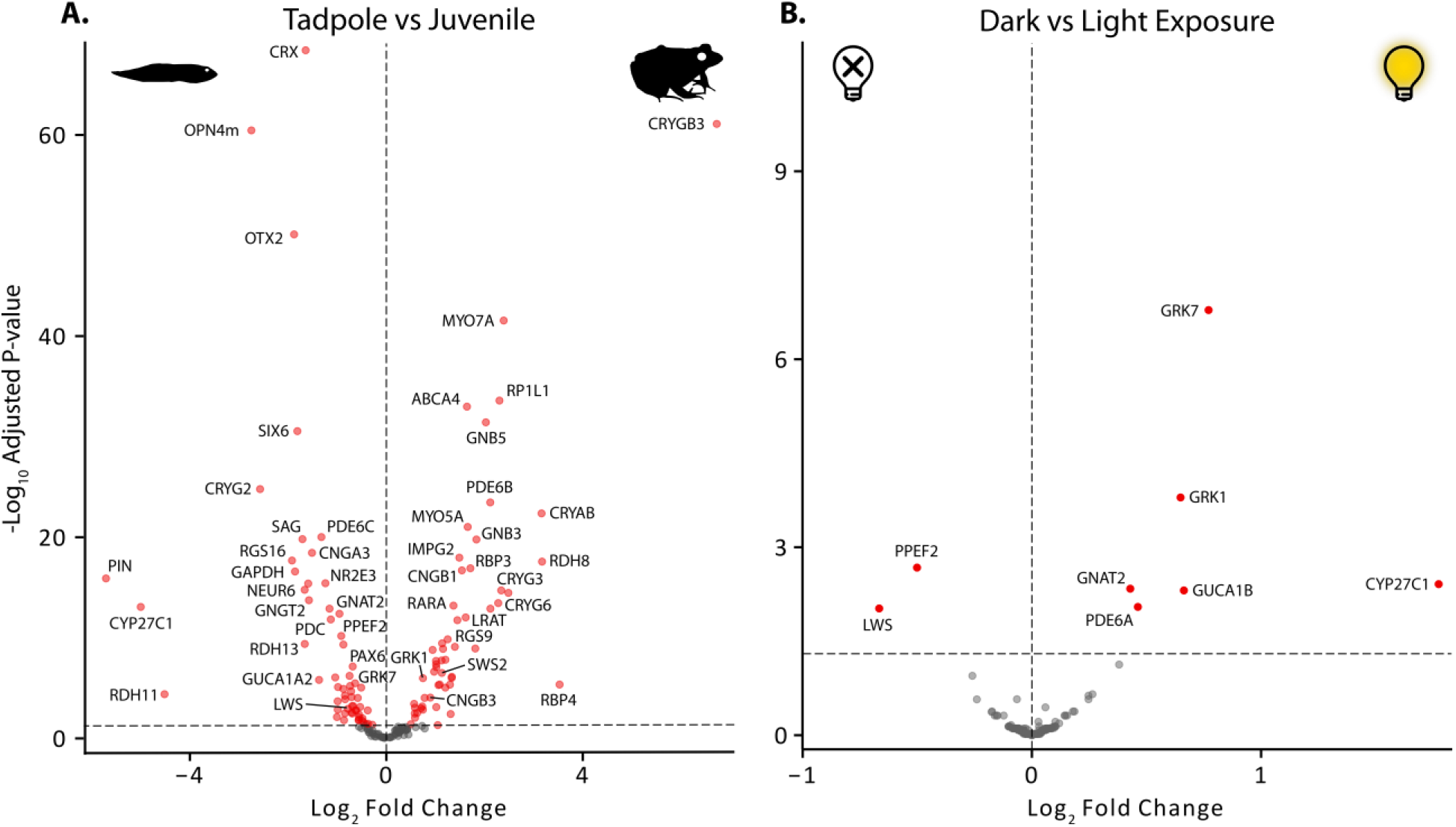
Volcano plots of differential expression of visual gene coding sequences between **A**) tadpoles and juveniles and **B**) dark and light exposure. Differential expression was estimated with DESeq2 using a multifactor design that accounted for the variation in both life stage (tadpole vs juvenile) and treatment (dark vs light exposure). For visualization, log_2_ fold changes (LFC) were shrunk and an adjusted p-value of 0.05 was set as the significance cut-off.

Eight genes (5%) were differentially expressed between light and dark-exposed individuals at an adjusted p-value of <0.05 (Fig. 3, Supplementary File 4), which is the same percentage of differentially expressed genes between light treatments in the transcriptome-wide analysis. These genes were a subset of those differentially expressed between tadpoles and adults, including several phototransduction genes, one visual opsin gene (*LWS*), and a gene involved in chromophore usage (*CYP27C1*, see below).

### *CYP27C1* is Upregulated in Tadpoles Coincident with a Red-Shift of the Spectral Absorbance of Visual Pigments

We found that *CYP27C1*, which encodes a cytochrome P450 family enzyme that converts A1 chromophore into A2 in vertebrates (Enright, et al. 2015), was significantly upregulated in tadpoles compared to juveniles, and in light exposed individuals (primarily tadpoles) compared to dark exposed (Fig. 4). Correspondingly, absorbance spectra of individual RH1 rod photoreceptor cells measured with MSP were red-shifted in the tadpole relative to the adult, and tended to fit models of A2 absorbance spectra better than A1 models (Fig. 4; Supplementary Table 3). Specifically, the absorbance maxima (λ_max_) of tadpole photoreceptors matched values expected from primarily A2-based visual pigments (e.g., 526 nm for RH1 rods), while λ_max_ values for adult photoreceptors matched expectations for primarily A1-based pigments (e.g., 505 nm for RH1 rods; Fig. 4; Supplementary Fig. 4). Estimation of λ_max_ and chromophore type was difficult for the other photoreceptor types due to the limited number of highly noisy scans we were able to obtain, especially for the tadpole due in part to its small size (Supplementary Fig. 5; Supplementary Table 2). However, the LWS cones appear to follow a similar pattern with a shift from A2-based pigment in tadpoles (λ_max_ of ~626 nm) to primarily A1-based pigment in adults (~579/603 nm; see below and Discussion for more details).

**Figure 4.**
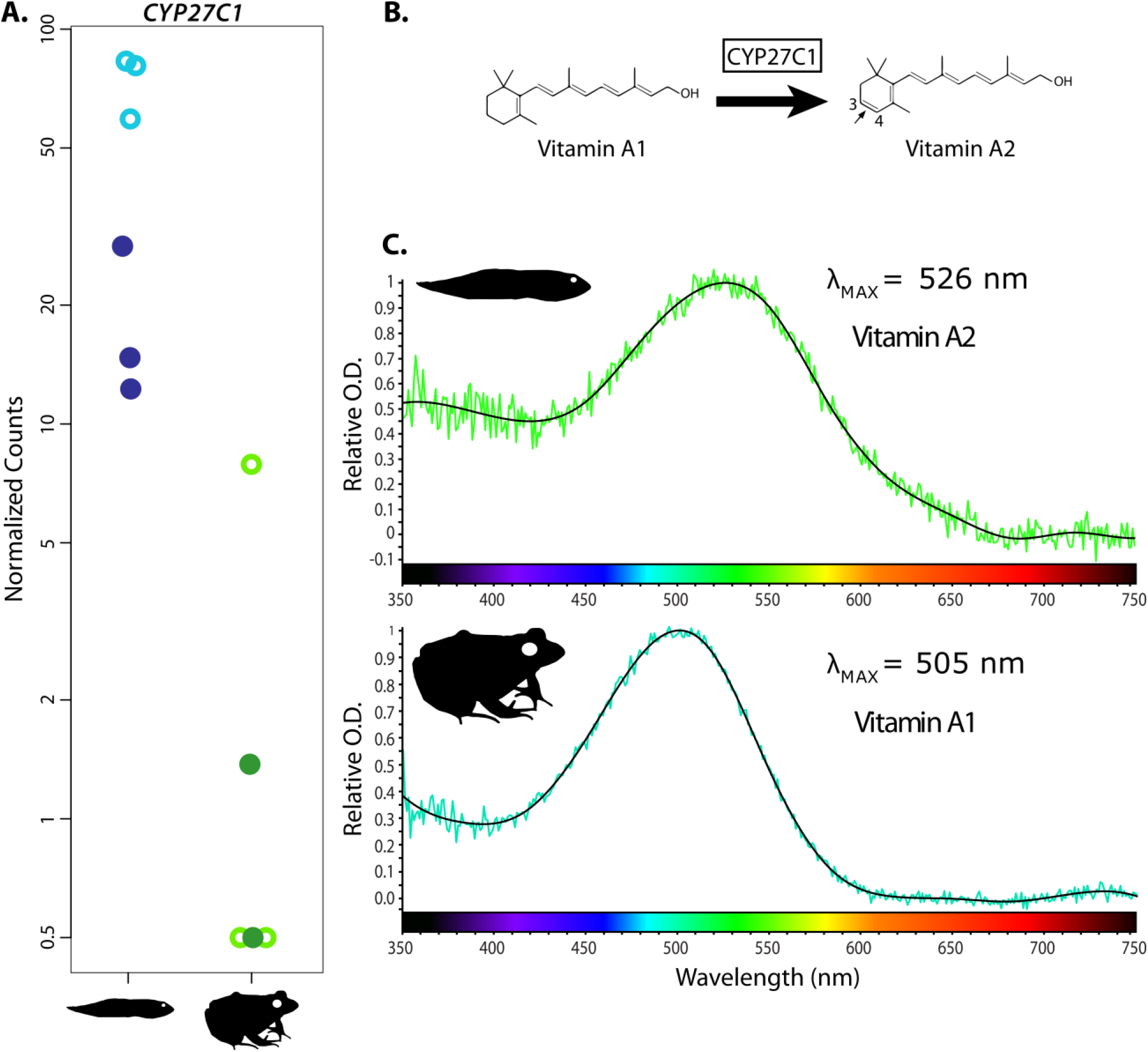
Upregulation of *CYP27C1* in tadpoles corresponds to red-shifted absorbance spectra of visual pigments and use of primarily A2 chromophore as detected by microspectrophotometry (MSP). **A**, Plots of normalized read counts of *CYP27C1* for light (open) and dark (closed) exposed juveniles and tadpoles. **B**, Conversion of vitamin A1 chromophore to vitamin A2 chromophore by *CYP27C1*. Based on Enright et al. (2015). **C**, Representative MSP absorbance spectra from RH1 (red) rod photoreceptors of an adult and a tadpole. Absorbance spectra for additional photoreceptor cell types and complete results tables can be found in Supplementary Figures 5–7; Supplementary Table 2, and Supplementary File 5.

### Three photoreceptor classes identified by MSP in tadpoles, with up to five classes in adults

We identified three photoreceptor classes in tadpoles using MSP: a blue-absorbing ‘green’ rod inferred to contain an SWS2 visual pigment with a λ_max_ of 433 nm (A1) / 431 nm (A2), a green-absorbing ‘red’ rod inferred to contain an A2-based RH1 pigment with a λ_max_ of 526 nm, and a red-absorbing cone inferred to have an LWS visual pigment with a λ_max_ of 635 nm (A1) / 626 nm (A2) (Supplementary Figs. 5–7; Supplementary Table 2, and Supplementary File 5; Schott et al. 2021). The SWS2 rod was from a single, very noisy scan and thus the λ_max_ and chromophore best fit (which was A1) may not be accurate. The LWS cone had an A1 chromophore best fit but the λ_max_ (635/626 nm) is inconsistent with an A1-based pigment (see Parry and Bowmaker (2000)). As such, we favoured the A2-based λ_max_ of 626 nm.

In adults we identified the same three photoreceptor classes as tadpoles, with the potential addition of a second RH1 rod type and a second LWS cone type. The SWS2 rod was inferred to contain an A1-based pigment with a λ_max_ of 437 nm, the primary RH1 rod an A1-based 505 nm pigment, with a secondary RH1 rod with a 505 (A1) / 501 (A2) nm pigment, and two LWS cones with A1-based 579 and 603 nm pigments, respectively (Supplementary Fig. 7; Supplementary Table 2). The secondary RH1 rod had an A2 chromophore best fit, but with λ_max_ of 501 nm an A2-based pigment is unlikely. Thus, it is unclear whether these truly represent two distinct photoreceptor populations (see Discussion).

### Visual Opsin Expression Varies between Life Stages and Light Treatments

Each of the four visual opsin genes expected in frogs (*RH1*, *LWS*, *SWS1*, and *SWS2*) were expressed in both tadpoles and juveniles. The rod opsin *RH1* was the most highly expressed of the four, likely reflecting the high number of RH1 rod photoreceptor cells typically found in frog retinas. Although our MSP approach is not appropriate for quantitative estimates of photoreceptor abundances, RH1 rod photoreceptors were the dominant cell type we observed and measured (Supplementary Table 2), which matches expectation based on studies in *L. pipiens* (Nilsson 1964; Liebman and Entine 1968). *RH1* expression showed high individual variability but did not differ with respect to life stage or light exposure (Fig. 5). The cone photoreceptor opsin genes *LWS* and *SWS2* had similar relative expression levels and we detected corresponding photoreceptor types for both opsins with MSP (Supplementary Fig. 5; Supplementary Table 2). *SWS2* was significantly upregulated in juveniles compared to tadpoles, while *LWS* was significantly upregulated with dark vs light exposure, although this was driven by high relative expression in tadpoles, specifically. Finally, the cone opsin gene *SWS1* showed the lowest relative expression, and though consistently expressed with some individual variation, showed no shifts associated with life stage or light exposure. Despite the expression of this gene, no short wavelength sensitive cones were detected with MSP (Supplementary File 5).

**Figure 5.**
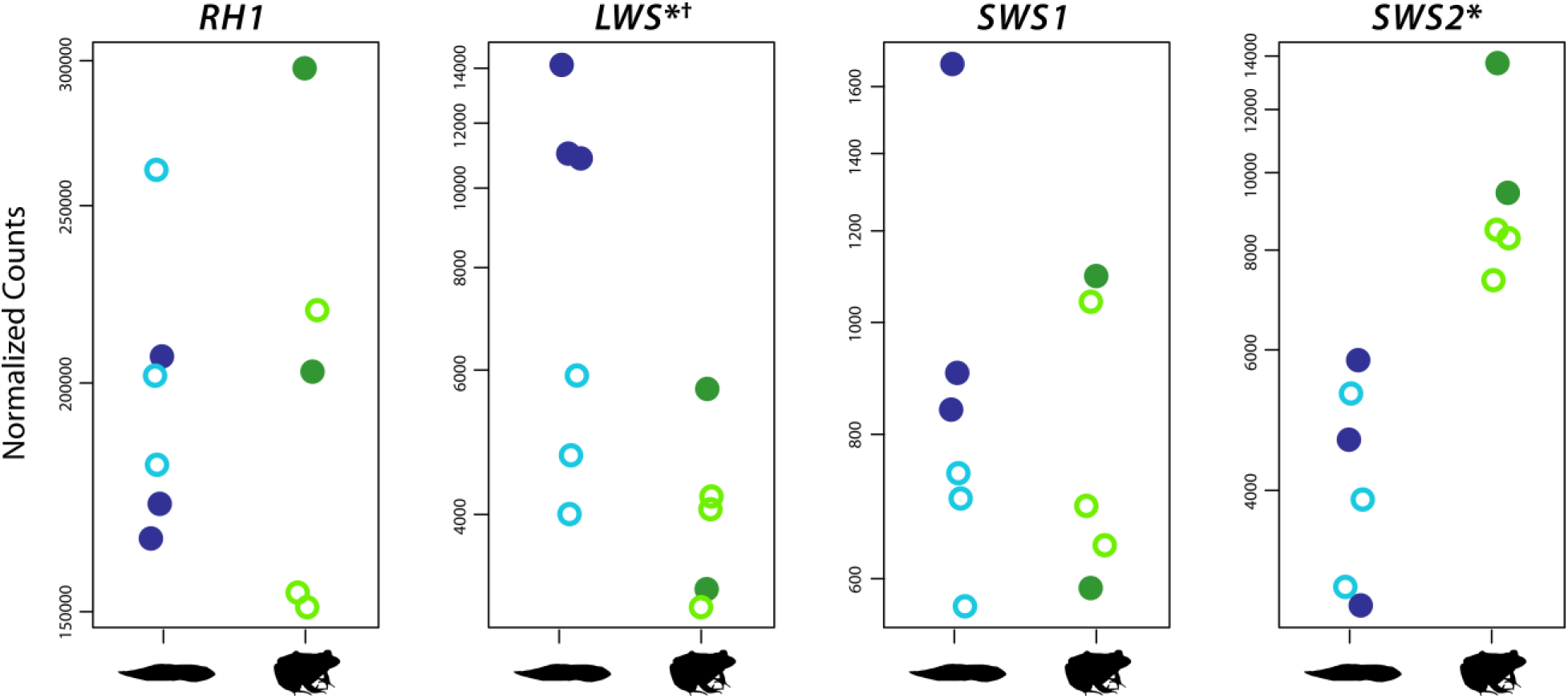
Expression profiles of visual opsin genes. *LWS* and *SWS2* were significantly differentially expressed between juveniles and tadpoles (indicated by the *), although the difference for *LWS* was driven by significant upregulation with dark exposure (indicated by the ^†^), specifically in tadpoles. Plots are of normalized read counts for each gene. Dark and light exposure are denoted by the closed and open circles, respectively. RH1, LWS, and SWS2 photoreceptor cell types were detected by MSP, whereas SWS1 was not (corresponding MSP results in Supplementary Figs. 5–7 and Supplementary Table 2).

### Expression Profiles of Lens Crystallin Genes Shift between Tadpoles and Juveniles

We analyzed gene expression patterns for 21 ubiquitous crystallins (two ɑ-crystallins, six β-crystallins, and 13 γ-crystallins) (Wistow 1995), and two ‘taxon-specific’ crystallins previously reported in frogs (ρ-crystallin and ζ-crystallin) (Tomarev, et al. 1984)

(Fujii, et al. 2001). Of these, two β-crystallin (*CRYBA1*, *CRYBB1*), two γ-crystallin (*CRYG2*, *CRYGN*), and both taxon-specific crystallin genes were significantly upregulated in tadpoles (Fig. 6, Supplementary File 4). One α-crystallin (*CRYAB*), two β-crystallin, and four γ-crystallin genes were significantly upregulated in juveniles. Overall changes in relative expression of α-, β-, and γ-crystallin genes were investigated using TMM cross-normalized expression values. Relative expression of γ-crystallin genes was more than twice that of α- and β-crystallin genes in tadpoles (691,387 vs 296,175; Supplementary Fig. 7; Supplementary Table 3) and slightly less than twice that in juveniles (591,814 vs 313,789). Relative expression of α-crystallin genes was higher in juveniles (96,040 vs 132,927), while β-crystallin gene expression was higher in tadpoles (200,135 vs 180,862). Compared to the ubiquitous crystallins, the two taxon-specific crystallin genes made up a relatively small proportion of the crystallin expression (3.8%) with ρ-crystallin comprising 99.9% of this expression. The relatively low expression of ζ-crystallin suggests it may not be a component of the lens in leopard frogs and may instead be present at housekeeping levels within the eye, because this protein has additional roles outside of the lens, at least in mammals (see Lapucci, et al. (2010) and references therein). Finally, we analyzed expression patterns of five other taxon-specific crystallins that have not specifically been identified in frog lenses in previous studies, and one (α-enolase/ENO1/τ-crystallin) with mixed evidence (Krishnan, et al. 2007; Keenan, et al. 2009). Of these, three were differentially expressed (two upregulated in tadpoles, one in juveniles), including α-enolase (Supplementary Fig. 8). We found no evidence for an effect of light exposure on the expression of any of the lens crystallin genes.

**Figure 6.**
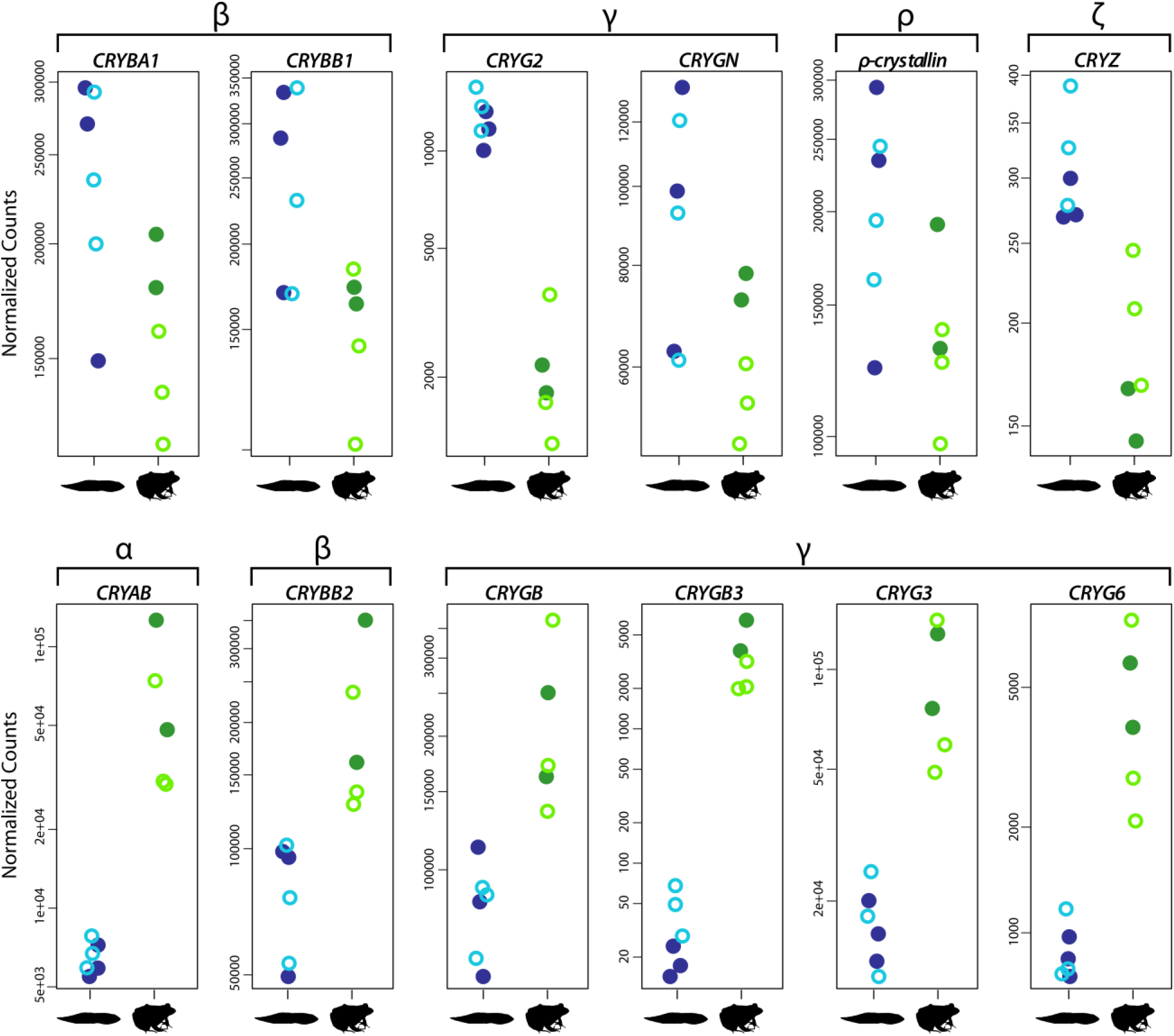
Expression profiles of lens crystallin genes differ substantially between tadpole and adults. Each of the crystallin genes depicted was significantly differentially expressed between juveniles and tadpoles (adjusted p-value < 0.05; Supplementary Table 3). The top row contains crystallins upregulated in tadpoles, which includes two β-crystallins (*CRYBA1*, *CRYBB1*), two γ-crystallins (*CRYG2*, *CRYGN*) and two taxon-specific crystallins (ρ-crystallin and ζ-crystallin, *CRYZ*). The bottom row contains those upregulated in juveniles, which includes one α-crystallin (CRYAB), one β-crystallin (*CRYBB2*), four γ-crystallins (*CRYGB*, *CRYGB3, CRYG3, CRYG6*). Plots are of normalized read counts for each gene with light and dark exposed samples denoted by the open and closed circles, respectively.

To explore potential functional consequences of changes to lens crystallin composition, we estimated protein refractive index increments (*dn*/*dc*) for the ubiquitous and known frog taxon-specific crystallins (Supplementary Fig. 9, Supplementary Table 3, Supplementary File 6). The *dn*/*dc* value defines how much a particular protein will contribute the refractive index at a given concentration (Mahendiran, et al. 2014). Of the ubiquitous crystallins, leopard frog α-crystallins were estimated to have the lowest *dn*/*dc* values (0.1941), followed by β-crystallins (0.1970), and γ-crystallins (0.2021). The α-crystallin that was significantly upregulated in juveniles (encoded by *CRYAB*) had the lowest *dn*/*dc* of any of the ubiquitous crystallins (0.1934). The two β-crystallins encoded by significantly upregulated genes in tadpoles had higher *dn*/*dc* values (CRYBA1, 0.1983; CRYBB1, 0.1952) than the β-crystallin upregulated in juveniles (0.1948). The pattern among the differentially expressed γ-crystallins was less clear. The two upregulated γ-crystallins in tadpoles both have relatively high *dn*/*dc* values, but the four upregulated in juveniles had a mix of values. The γ-crystallins, in particular, also had highly variable expression levels and therefore would likely contribute very different amounts to the overall refractive index. When accounting for expression levels (TMM) of all the ubiquitous crystallin, tadpoles had slightly higher average (weighted mean) *dn*/*dc* than juveniles (0.2010 vs 0.2003; Supplementary File 6). The frog-specific crystallins had lower estimated *dn*/*dc* values than any of the ubiquitous crystallins (ρ-crystallin, 0.1916 and ζ-crystallin, 0.1884; Supplementary Table 3).

## Discussion

### Significant Shifts in Gene Expression across Ontogeny

The adaptive decoupling hypothesis suggests that animals that develop through two or more distinct phenotypic phases may be able to disrupt genetic correlations between phases, allowing each phase to better adapt to its particular selective environment. This strategy would be particularly relevant for frogs, which often have larval (tadpole) and adult phases that occupy very different ecological niches. Although previous studies have shown that morphological diversity of tadpoles and adult frogs is decoupled (Roelants, et al. 2011; Sherratt, et al. 2017; Wollenberg Valero, et al. 2017; Fabrezi, et al. 2019), studies testing for differences in gene expression are more limited (Wollenberg Valero, et al. 2017). We found that a large proportion of the eye transcriptome (42%) was significantly differentially expressed between tadpole and juvenile southern leopard frogs, suggesting substantial, and potentially adaptive, decoupling at the level of gene expression. These results are consistent with a recent study of eye-body size allometry across ontogeny in anurans that detected a shift at metamorphosis in several species (Shrimpton et al. *in press*), suggesting that relative eye growth is partially decoupled as well. The extent of decoupling in gene expression we detected was substantially more than was found in a comparison of tadpole and adult *Mantidactylus betsileanus* Madagascar frogs whole body (excluding gut) transcriptomes, which found that only 14% of annotated transcripts were differentially expressed (Wollenberg Valero, et al. 2017). In that whole body comparison, differentially expressed transcripts were significantly enriched for genes involved in morphological development, suggesting that phenotypic evolution across phases was decoupled.

Other examples of decoupling in gene expression come from insects (Jacobs, et al. 2006; Fellous and Lazzaro 2011; Saenko, et al. 2012; Herrig, et al. 2020). For instance, in the hymenopteran *Neodiprion lecontei*, which has multiple life stages separated by increasingly dramatic metamorphic transitions, Herrig et al (2020) found that a progressively greater proportion of genes (up to 31%) were differentially expressed between each successive pair of stages. That observation matched the authors’ predictions that gene expression decoupling would be strongest between the most ecologically dissimilar life stages. Consequently, we propose that expression may be more strongly coupled in frog species that remain aquatic after metamorphosis or in those that do not have a tadpole life stage and instead complete development within the egg and hatch as juvenile frogs (direct development). Further studies with additional species and finer scale ontogenetic sampling are needed to test this hypothesis.

Although our results have interesting implications for the broad decoupling of gene expression between tadpoles and juveniles, our focus was on visual genes. We found that the visual system was disproportionately decoupled compared to the full eye transcriptome, with significant differential expression between tadpole and juvenile frogs in over half of the visual genes we analyzed. This suggests the visual system may be particularly strongly adapted, through decoupling, to the larval and adult environments. Teleost fishes also show differences in visual gene expression across ontogeny, specifically with regards to visual opsin expression, when larval habitat or foraging differs from that of adults (Carleton, et al. 2020). Changes in expression patterns of other visual genes in teleosts remain to be evaluated, but we expect considerable variation in those taxa with distinct larval and adult ecologies. Likewise, it has recently been proposed that the visual system of the tuatara (*Sphenodon punctatus*) may be uniquely adapted to accommodate differing juvenile and adult ecologies due to the constraints imposed by a single life stage (Gemmell, et al. 2020). Tuatara juveniles often take up a diurnal and arboreal lifestyle to avoid the terrestrial, nocturnal, adults that may predate them, and this appears to have resulted in a visual system with a unique mix of diurnal and nocturnal features and highly conserved in visual genes compared to other vertebrates (Gemmell, et al. 2020). Consequently, gene expression decoupling across ontogeny may also feature in tuatara despite the absence of a complex life cycle. Overall, these studies suggest that differential expression of visual genes may be a fairly widespread strategy in vertebrates that adapt to different light environments across ontogeny.

The visual genes we identified as differentially expressed between tadpole and juvenile leopard frogs effect a broad range of visual functions, including developmental genes, such as *CRX* and *OTX2,* that are important for the differentiation and maintenance of rod and cone photoreceptor cells (Yamamoto, et al. 2020). Several non-visual opsins were also differentially expressed, including melanopsin (*OPN4m*) and pinopsin (*PIN*), suggesting that circadian clocks and regulation may differ between tadpoles and juveniles (Poletini, et al. 2015). A number of phototransduction and visual cycle genes were differentially expressed, which is likely to have further enabled the partially independent adaptation of the leopard frog visual system to the larval and adult environments. Some of these genes were also differentially expressed between the light and dark exposure treatments, which may reflect diel expression patterns or an ability to plastically acclimate to different light environments at the level of gene expression. Finally, the chromophore usage, visual opsin, and lens crystallin genes also had strong support for differential expression between tadpoles and juveniles and have clear consequences for vision in different light environments, which are discussed in more detail below.

### Significant Shifts in Gene Expression with Light Exposure

In addition to differential gene expression across ontogeny we also found a smaller set of genes (5% of total) were differentially expressed between dark and light exposed treatments, which was consistent across the full transcriptome and the visual gene datasets. This experimental design was primarily motivated by ongoing comparative studies by our group, and others, that use eye and retinal transcriptomes for molecular evolutionary analyses. In many cases precise control of lighting environments is not possible (e.g., field collected samples) and we aimed to explore the potential impacts of light exposure variation on visual gene recovery from transcriptomes. Our results suggest that variation in light conditions during samping is unlikely to have a significant impact on comparative studies that aim to recover visual genes from whole-eye transcriptomes and conduct molecular evolutionary analyses. However, we did find that several phototransduction genes (e.g., rod and cone opsin kinase [*GRK1* and *GRK7*], cone transducin [*GNAT2*]), one visual opsin (*LWS*), and the chromophore usage gene (*CYP27C1*) were significantly differentially expressed between light and dark exposure. Consequently, we infer that controlled light environments and sampling conditions are likely necessary for accurately assessing differential expression of visual genes in comparative studies. In addition, we suggest that variation in gene expression between light and dark treatments may represent a form of adaptive plasticity to short-term changes in light levels that warrants further investigation both within frogs and in other vertebrates.

### Chromophore Usage and Light Sensitivity Shift Across Ontogeny

Changes in visual pigment light sensitivity associated with the use of the A2 and A1 chromophore have been established using spectrophotometry of retinal extracts and MSP of intact photoreceptors in several tadpole and adult frogs (reviewed in Bridges 1972). Recently the gene responsible for converting A1 into A2, *CYP27C1*, was identified (Enright et al. 2015). We found that *CYP27C1* was significantly upregulated in southern leopard frog tadpoles compared to juveniles and, with MSP, confirmed that this difference is associated with red-shifted absorbance spectra and A2 chromophore usage in tadpole RH1 rods. Thus, our differential expression and MSP results provide evidence for the role of *CYP27C1* in switching chromophore usage and, correspondingly, spectral sensitivity during ontogeny. Several species of frog that transition from aquatic tadpoles to terrestrial or semi-terrestrial adults, including the Pacific tree frog *Hyla regilla* and several other *Lithobates* species (*L. catesbeianus*, *L. clamitans*, *L. esculenta*, *L. pipiens, L. sphenocephalus*, *L. temporaria*), shift their chromophore usage from primarily A2 to primarily A1 at metamorphosis (Muntz and Reuter 1966; Liebman and Entine 1968; Bridges 1972, 1974). By contrast, the toads *Bufo boreas* and *B. bufo* appear to use only A1 as both aquatic tadpoles and terrestrial adults (Crescitelli 1958; Muntz and Reuter 1966), and *X. laevis*, which remain fully aquatic as adults, use exclusively A2 pigments at both life stages (Bridges, et al. 1977). Additional studies quantifying ontogenetic variation in chromophore usage and *CYP27C1* expression across a greater diversity of frogs may provide a better understanding of how this trait is associated with changes in visual ecology across species.

Light exposure can also affect the proportion of A1/A2 chromophores in the eye. This pattern has been demonstrated as a reversible change in tadpoles of several species of *Lithobates,* where exposure to darkness over extended periods (weeks to over a month) decreased the proportion of A2 chromophore, which could be reversed to near normal levels in as little as 24–48h of constant illumination (Bridges 1970, 1974). We found that *CYP27C1* expression was significantly reduced in dark-exposed compared to light-exposed tadpoles (12h of exposure), which suggests *CYP27C1* expression is light-dependent in tadpoles and that reduced *CYP27C1* expression upon dark exposure results in a lower proportion of A2-based visual pigments. A potential mechanism for light-dependent expression of *CYP27C1* is unknown, but previous experiments suggest this response can be localized to a single eye. Specifically, Bridges (1974) found that exposing tadpoles to light after a period of darkness resulted in an increase in A2 in unobstructed eyes, but not in eyes that were covered and thus unexposed to light. The functional benefit of this change may be related to the lower dark noise (thermal activations in the absence of photon absorbance) of A1 pigments compared to A2 pigments, resulting in higher light sensitivity of A1 pigments (Ala-Laurila, et al. 2007). Thus, exposure to dark conditions could result in increased overall light sensitivity of the tadpole through reduced expression of *CYP27C1*, which in turn reduces the production of A2 chromophore and, correspondingly, increases the proportion of the higher-sensitivity A1 pigments. This interpretation is perhaps confounded by the observation that some teleost fish species have the opposite reaction to dark exposure, where the proportion of A2 pigment increases (Bridges 1972), although at least one teleost species has the same response as tadpoles (Allen 1971). This variation suggests there may be other factors beyond spectral tuning and light sensitivity that influence chromophore usage.

### Visual Opsins are Differentially Expressed Between Life Stages

Differential expression of visual opsins is a second approach that organisms can employ to adapt the sensitivity of their visual system to changes in the light environment. Studies investigating the evolutionary and ecological context of this particular strategy in vertebrates have largely been restricted to teleost fishes, which tend to have many more (duplicated) copies of visual opsins than do tetrapods (Carleton, et al. 2020). Some teleost species express only a subset of their total visual opsin repertoire during a particular life stage, which is likely beneficial when larval habitat and associated traits, such as foraging behavior, differ from those of adults (Carleton, et al. 2020). Most frog species make even more dramatic transitions in habitat and foraging behavior across ontogeny, but with far fewer opsins to choose from, their ability to use distinct opsins for larval and adult vision is likely more constrained. Despite this, we found differential expression between tadpoles and adults in two of their four visual opsin genes (*SWS2* and *LWS*). In frogs and salamanders, *SWS2* is expressed in blue absorbing ‘green’ rods, a novel class of rod photoreceptor that, at least in some species, enables colour vision at scotopic light levels when cones are inactive (Yovanovich, et al. 2017). The increased expression of *SWS2* in juvenile leopard frogs suggests an increased proportion of SWS2 rods in the retinas of juveniles relative to tadpoles, which may reflect increased reliance on nocturnal colour vision in terrestrial life stages. By contrast, the significant differences in *LWS* expression appears to be driven by the effect of dark exposure in tadpoles. Previous studies in teleost fishes have found variable responses of opsin expression in different light conditions (Carleton, et al. 2020; Cortesi, et al. 2020) including expression changes with respect to housing animals in different light environments (e.g., clear vs tea-stained water; (Fuller, et al. 2010; Fuller and Claricoates 2011)) and daily (diel) variation in expression (Stieb, et al. 2016; Yourick, et al. 2019). One common observation in daily expression cycles is that cone opsin expression peaks near the onset of darkness and remains high throughout the night (Halstenberg, et al. 2005; Yourick, et al. 2019). Such variation has not yet been characterized in anurans, and future studies with broader taxonomic sampling, and a greater range of light/dark treatments, would improve our understanding of visual gene expression plasticity in larval and adult anurans.

Although we found *SWS1* opsin gene expression in both tadpole and adult life stages, we did not find evidence for any short-wavelength sensitive (SWS) cone photoreceptors that could be attributed to either of the short-wavelength sensitive opsins (SWS1, SWS2) with MSP. Relative expression of *SWS1* was the lowest of the four visual opsins, suggesting that if SWS1 photoreceptors exist in leopard frogs, they may be rare and/or small and thus difficult to detect and measure with MSP. In both *X. laevis* and *L. catesbeianus*, SWS1 opsin was localized to a subset of single cone photoreceptors using immunohistochemistry, while SWS2 was exclusive to ‘green’ rods (Hisatomi, et al. 1998)Hisatomi, 1999 #1163}(Starace and Knox 1998; Darden, et al. 2003). This is in contrast to salamanders, which can have both SWS1 and SWS2 cones in addition to SWS2 ‘green’ rods (Ma, et al. 2001; Isayama, et al. 2014). Using MSP, *L. catesbeianus* was found to have blue-sensitive cones with the same λ_max_ (433 nm) as the SWS2 ‘green’ rods (Hárosi 1982), which when combined with the immunohistochemistry data (Hisatomi, et al. 1998; Hisatomi, et al. 1999) suggests that the SWS1 and SWS2 visual pigments may have converged on the same absorbance spectrum in this species (Donner and Yovanovich 2020). Blue-sensitive cones have not been found with MSP in *X. laevis* (Witkovsky, et al. 1981), but the absorbance spectra of the SWS1 and SWS2 visual pigments were measured *in vitro* (with the A1 chromophore) and found to differ by almost 10 nm (425 and 434 nm, respectively). Thus, it is possible that frogs co-express SWS1 with other visual pigments, as occurs in salamanders and several other vertebrate groups (Dalton, et al. 2014; Isayama, et al. 2014). This could also explain the difficulty in detecting SWS1 in southern leopard frogs with MSP. Additional studies are clearly needed to further resolve the photoreceptor complements of frogs and how they vary across taxa.

### Photoreceptor Spectral Sensitivity Variation in Closely Related Leopard Frog Species

Our *Lithobates sphenocephalus* MSP results are fairly similar to spectral sensitivities previously reported for the closely-related northern leopard frog (*L. pipiens*; Liebman and Entine (1968); Liebman (1972)) but with some notable differences. Estimates for RH1 rods across species only differed by 1 nm in tadpoles (λ_max_ = 526 vs 527 nm [A2] for *L. sphenocephalus* and *L. pipiens*, respectively). The protein sequences for RH1 are nearly identical between the two species and so the resulting visual pigments are expected to have very similar absorbances. In adult *L. sphenocephalus*, we were able to distinguish two classes of rods: the more numerous class had a λ_max_ of 505 nm with an A1 best fit, while the less numerous had a λ_max_ of 501 nm with an A2 best fit. Interestingly, Liebman and Entine (1968) also classified two RH1 rods in adult *L. pipiens*: the more numerous with λ_max_ at 502 nm and a rarer class with λ_max_ as high as 507 nm, both thought to be A1-based. The higher λ_max_ of some rods could be due to a mixture of a small amount of A2-based pigment with the A1-based pigment in a subset of rod cells. Liebman and Entine (1968) argued against this possibility because they found both rod types had the same A1 bleaching intermediates in *L. pipiens*, but it is unclear what other mechanism could account for this observation. Alternatively, the two RH1 rod classes could be due to the presence of two RH1 alleles that encode spectrally distinct RH1 opsin proteins, but we found no evidence to support additional RH1 alleles based on our *de novo* assembled eye transcriptome data. Thus, we tentatively conclude the second RH1 rod class in our MSP dataset is most likely due to a small proportion of A2-based pigment, which is further supported by the variable levels of *CYP27C1* expression we found for the juvenile individuals. The presence of two RH1 rod classes could be related to spatial variation in the ratio of A1- to A2-based visual pigments across the retina as found in adult bullfrogs (*L. catesbeianus;* Enright, et al. (2015)). Quantitative studies of chromophore content would more conclusively differentiate among these potential explanations.

Changes in spectral sensitivity of SWS2 rods showed opposite trends across life stages in southern vs northern leopard frogs, with higher λ_max_ in adults of *L. sphenocephalus* (437 vs 433 nm in tadpoles) versus higher in tadpoles of *L. pipiens* (438 vs 432 nm in adults). Unfortunately, our SWS2 rod estimate from the *L. sphenocephalus* tadpole was based on only one noisy scan, so our estimates of λ_max_ and chromophore type may be inaccurate. However, our results do support a small (~5 nm) red-shift for the adult SWS2 rods in *L. sphenocephalus* relative to *L. pipiens*. For the tadpole LWS cones, we found a similar red-shift between species (λ_max_ = 626 nm in *L. sphenocephalus* vs 620 nm [A2] in *L. pipiens*). In adults, the 579 nm cones we identified in *L. sphenocephalus* are similar to the A1-based 575 nm LWS cones reported for *L. pipiens* (Liebman and Entine 1968), but again slightly red-shifted. This implies that the second type of LWS cone we found (603 nm) is the result of a small amount of A2 pigment mixed with A1-based pigment, also resulting in a red shift. Unfortunately, protein sequences for *L. pipiens* SWS2 and LWS are not available for comparison to determine if changes to the protein sequence are likely to contribute to the observed differences in λ_max_. Overall, these results suggest there may be a large amount of unappreciated variation in frog photoreceptor complements and spectral sensitivities.

### Lens Crystallin Expression and Estimated Refractive Index of the Lens Shifts across Ontogeny

Previous studies of frog lens crystallin proteins found predominantly γ-crystallins (CRYG) in tadpoles and α- and β-crystallins in adults (CRYA and CRYB) (Polansky and Bennett 1973; Doyle and Maclean 1978; Hoskins 1990), which is partly consistent with our observations of crystallin gene expression. Although the combined level of α- and β-crystallins increased relative to γ-crystallins in juveniles, β-crystallin expression, specifically, decreased slightly, which is contrary to expectations based on protein studies in *L. catesbeianus* (Polansky and Bennett 1973; Jiang, et al. 1989). However, it is possible that with further growth a similar shift would be observed in β-crystallin gene expression in *L. sphenocephalus* considering that changes in crystallin composition have been linked to increases in lens diameter rather than with metamorphosis (Doyle and Maclean 1978). Beyond the overall changes in α-, β-, and γ-crystallin expression, our findings suggest a more complicated scenario of crystallin usage at the gene level. Different β- and γ-crystallin genes were upregulated in tadpoles versus juveniles indicating turnover in both of these types of crystallins.

One of the primary roles of crystallins is to provide a high refractive index to the lens (Zhao, Brown, Magone, et al. 2011). Experimental and computational studies have estimated refractive index increments for multiple crystallins, and α-crystallins were found to have the lowest refractive indices, followed by β-, and then γ-crystallins, which have exceptionally high values compared to other proteins (Pierscionek, et al. 1987; Zhao, Brown, Magone, et al. 2011). Our estimates of refractive increments for the leopard frog ubiquitous crystallins agreed with this general trend. Consequently, we predict that the increased expression of *CRYAB* (the encoded crystallin of which had the lowest estimated refractive index) and decreased expression of γ-crystallins in juveniles would reduce the refractive index of their lens relative to tadpoles. This change in refractive index across ontogeny likely reflects the need to avoid overfocusing (myopia/nearsightedness) when juveniles transition to vision in air, where the cornea provides substantial refractive power. This explanation is further supported by Mahendiran, et al. (2014) who found aquatic vertebrates (*X. laevis* and zebrafish) generally had crystallins with higher refractive increments than terrestrial mammals.

We found that the two β-crystallins encoded by genes upregulated in tadpoles (CRYBA1, CRYBB1) had higher refractive increments than the β-crystallin gene upregulated in juveniles (CRYBB2), suggesting that a similar shift in the refractive increments of crystallins could contribute to a change in the refractive index of the lens across ontogeny in frogs with aquatic larval and terrestrial adult life stages. However, it should be noted that the pattern within γ-crystallins was not as clear, and that when accounting for relative expression levels (as a rough approximation for protein abundances) we found that the average refractive increment of the crystallin composition in juveniles was only slightly less than that in tadpoles. It has been proposed that the shift in crystallin composition from primarily γ-crystallin to α- and β-crystallins serves to maintain the refractive index of the frog lens during growth, while an increased hydration of the lens, along with the change in lens shape, may be responsible for the decreased power of the lens in the transition to vision in air (Smith-Gill and Carver 1981). The difference in γ-crystallin usage could also be related to a change in lens hydration. Zhao, et al. (2014) found that zebrafish γ-crystallins exhibited extremely low degrees of hydration consistent with their role in high refractive index aquatic lenses. This was contrasted with average and low degrees of hydration of different mouse and human γ-crystallins. A shift in γ-crystallin usage in frogs that transition from aquatic to terrestrial habitats could facilitate a change in lens hydration, but this remains to be tested. However, these results highlight that additional studies are needed both to examine frog lens crystallins more directly, and to examine how they turnover across the full range of ontogeny, to better understand how the lens has adapted in response to different light environments.

Two taxon-specific crystallins have been identified in frog lenses, but the specific roles they play in the function of the lens have not been studied. ρ-crystallin (originally referred to as ε-crysallin) was first found in *Rana temporaria* and later in *L. catesbeianus* (Tomarev, et al. 1984; Fujii, et al. 1990), while ζ-crystallin was identified in the lenses of the hylid frogs *Hyla japonica, Litoria infrafrenata,* and *Phyllomedusa sauvagei* (Fujii, et al. 1990; Keenan, et al. 2012). We found significant differential expression of both ρ- and ζ-crystallin genes in leopard frog eyes; however, the very low relative expression of ζ-crystallin suggests that it does not function as a lens crystallin in this species. A third taxon-specific crystallin, α-enolase (ENO1, τ-crystallin) has mixed reports regarding its presence in the lenses of *Bufo* toads (Krishnan, et al. 2007; Keenan, et al. 2009). We found that *ENO1* was differentially expressed across life stages and expressed at moderate levels, but like most other taxon-specific crystallins, ENO1 has varied enzymatic functions and broad tissue expression (Ji, et al. 2016), thus its role as a lens crystallin in leopard frogs requires further investigation.

## Conclusions

We found high levels of decoupling of gene expression between aquatic tadpole and terrestrial juvenile southern leopard frogs consistent with the adaptive decoupling hypothesis. The degree of decoupling was even greater among visual genes, suggesting that adaptive decoupling may have played an important role in the evolution and adaptation of anuran visual systems. Specifically, our results highlight expression differences in a range of visual genes, including chromophore usage, visual opsin, and lens crystallin genes, that likely underlie observed morphological and physiological changes through metamorphosis and corresponding adaptive shifts to optimize visual ability in aquatic versus terrestrial light environments. We also found evidence that light/dark exposure has a significant effect on the expression of a much smaller, but similar, set of visual genes. These results set the stage for investigating adaptive decoupling and differential expression of visual genes across a broader ecological sampling of larval and metamorphosed anurans and further investigating the plasticity of visual gene expression in vertebrates.

## Materials and Methods

### Study Animals

Twelve southern leopard frogs (*Lithobates sphenocephalus* [=*Rana* (*Pantherana*) *sphenocephala*, (Yuan, et al. 2016)]; six tadpoles and six juveniles) were obtained from a single wild population in Arlington, TX in May and June 2018 (*-*97.101168, 32.792202; Texas Parks and Wildlife Scientific Research Permit No. SPR-0814-159; UTA IACUC Protocol A17.005) (Supplementary Table S1). The tadpoles were Gosner stages 25–38 and juvenile snout-vent lengths (SVL) were 24.77 mm to 34.59 mm (less than 51 mm, which is the minimum SVL leopard frogs are reported to reach sexual maturity at (Lannoo 2005). Three tadpole and three juvenile specimens were exposed to ambient light in the laboratory or complete darkness (ie., dark adapted) for 12 hours prior to euthanasia (via imersion in a solution of MS222). One whole eye from each specimen was extracted and placed in RNAlater (Ambion) for at least 24 hours at 4°C to allow the RNALater to saturate the cells, prior to freezing and storage at −80°C until use. For the dark-adapted specimens, eyes were dissected in the dark, placed in RNAlater, and then wrapped in foil to keep them in the dark during the entire process. An additional tadpole (not staged) and an adult (SVL 66.6 mm) were collected in April and June 2019 from the same location for microspectrophotometry (MSP). Voucher specimens are accessioned at the Amphibian and Reptile Diversity Research Center at UT Arlington and Smithsonian Institution’s National Museum of Natural History (Supplementary Table 1).

### Transcriptome Sequencing and Assembly

Total RNA was extracted from whole eyes using the Promega Total SV RNA Extraction kit (Promega). Tissue was homogenized in the prepared lysis buffer using the Qiagen Tissuelyzer (10 minutes at 20 Hz). Messenger RNA library construction was performed using the Kapa HyperPrep mRNA Stranded with Riboerase kit (Roche). Each indexed sample was pooled in equimolar amounts and sequenced on one lane of HiSeq4000 by Novogene.

Several quality control steps were employed to improve the quality and efficiency of the transcriptome assembly. Erroneous k-mers were removed from raw reads with rCorrector (Song and Florea 2015) and a custom script (available at https://github.com/harvardinformatics/TranscriptomeAssemblyTools). Adapters and low quality bases (q<5) were removed with TrimGalore! (https://www.bioinformatics.babraham.ac.uk/projects/trim_galore/), which implements Cutadapt (Martin 2011). Read pairs shorter than 36bp after trimming were discarded, as were unpaired reads. Ribosomal RNA reads were removed by mapping reads with Bowtie2 (Langmead and Salzberg 2012) against the SILVA database (Quast, et al. 2012). Quality of processed reads was assessed with FastQC (http://www.bioinformatics.babraham.ac.uk/projects/fastqc/). A *de novo* reference transcriptome was assembled using Trinity (Grabherr, et al. 2011) incorporating all paired reads from each of the 12 samples following the standard protocol. This assembly was subsequently redone removing one of the samples that was found to be an outlier in a principal component analysis of differential expression (see below). Alignment summary metrics were calculated using Trinity (Haas, et al. 2013). Read support for the assembly was determined by mapping the reads back to the assembly using Bowtie2 and completeness was assessed using BUSCO v3.0.2 (Seppey, et al. 2019) with the Tetrapoda dataset. The *de novo* Trinity assembly was reduced to a “best” set of transcripts using a modified version of the ‘Trinity best transcript set’ pipeline (https://github.com/trinityrnaseq/trinity_community_codebase/wiki/Trinity-best-transcript-set) available at (https://github.com/ryankschott/Best_Transcript_Set_Updated).

### Differential Expression and Gene Ontology Enrichment Analyses

Abundances of the reduced transcript set were quantified using Salmon (Patro, et al. 2017) and scripts included with Trinity. Differential expression was estimated with DESeq2 (Love, et al. 2014) using a generalized linear model with a negative binomial distribution and a multifactor design accounting for both life stages (tadpole vs juvenile) and treatment (light vs dark exposure). An adjusted p-value (padj) of <0.05 was used as the significance cutoff for differential expression. For data visualization and clustering, raw counts were transformed using the regularized logarithm (rlog), and log_2_ fold changes (LFC) were shrunk using the apeglm method (Zhu, et al. 2019). A principal component analysis (PCA) of rlog transformed counts was used to evaluate variation among the samples and to identify potential outliers. One of the juvenile, dark-exposed samples was identified as an outlier and removed from further analyses (Supplementary Fig. 1). Transcripts were annotated using BLASTn against the NCBI nucleotide (nt) database. To assess changes specific to the visual system, we generated a second dataset for differential expression analyses using a curated set of 170 visual gene coding sequences (Supplementary File 3). Reads were quantified against this reference using Salmon, and differential expression was estimated following the same approach outlined above. The same juvenile, dark-exposed sample was also identified as an outlier in the visual gene dataset and excluded from subsequent analyses (Supplementary Fig. 3). To compare relative expression among groups of genes, we used the Trinity pipeline to calculate cross-sample normalized (TMM) expression values (Robinson and Oshlack 2010).

To obtain gene ontology (GO) terms, we annotated the reduced set of transcripts using DIAMOND (Buchfink, et al. 2015) against the *Xenopus tropicalis* (v9.1) ENSEMBL database. GO enrichment analyses were performed using TopGO for the biological process gene ontology with the combined gene elimination and weighting algorithm (weight01) and Kolmogorov-Smirnov testing (KS) (Alexa and Rahnenfuhrer 2020).

### Microspectrophotometry

Microspectrophotometry was conducted on eyes from one tadpole and one adult (see above) following protocols described in previous studies (Loew 1994; Loew, et al. 2002). After at least 2h of dark adaptation, animals were euthanized with MS-222 and the eyes enucleated under dim red light. Eyes were hemisected, the cornea and lens isolated and the retinas carefully removed from the pigment epithelium under hypertonic buffer (pH 7.2 supplemented with 6% sucrose). Pieces of retina were macerated, sandwiched between two coverslips edged with silicone grease, and placed on the stage of a computer-controlled single-beam MSP (Loew 1994). Absorbance spectra were obtained for all clearly identified outer segments from 750 to 350 nm, and back again from 350 to 750 nm, with a wavelength accuracy of ~1 nm (Loew, 1994). Visual pigment λ_max_ was determined by template fitting using previously described methods (Loew 1994). Briefly, a Gaussian function was fit to the top 40 data points at 1 nm intervals and differentiated to establish the peak wavelength. The spectrum was normalised to this absorbance value and template fit to either A1 or A2 standard data (Govardovskii, et al. 2000) using the method of (MacNichol 1986). Template fitting alone is not the best determinant of A1 or A2 status for noisy data such as those from the very small outer segments of amphibian tadpoles and adult cones. However, if the calculated λ_max_ was greater than 580 nm, it was assumed that A2 must be present (Parry and Bowmaker 2000). Calculated λ_max_ values are accurate to ±1.0 nm and are reported here to the nearest whole integer.

### Protein Refractive Index Increment Estimation

Protein refractive index increments (*dn*/*dc*) were estimated for the predicted leopard frog lens crystallin proteins using the method of Zhao, Brown and Schuck (2011) as implemented in the SEDFIT software (Schuck 2010). The protein refractive index increment defines how much a given concentration of a protein contributes to the overall refractive index of the solution, which in the case of lens crystallin proteins equates to how much they will contribute to the refractive index of the lens. The method of Zhao and colleagues (2011) uses a biophysical computational model to estimate refractive index increments based on the amino acid composition of the protein and the refractivities of those amino acids as calculated by McMeekin, et al. (1962). Estimates were made assuming 589 nm light at 25°C and a solvent refractive index of 1.3340 following the protocol established by Zhao, Brown and Schuck (2011).

## Acknowledgements

Portions of the computational analyses were conducted on the Smithsonian Institution High Performance Cluster (SI/HPC; https://doi.org/10.25572/SIHPC). This work was supported by grants from the Natural Environment Research Council, UK (NE/R002150/1) and the National Science Foundation, USA (DEB- #1655751). We thank Nihar Bhattacharya for discussions on A2 visual pigments, and Corey Roelke for assistance in the field.

## Data Availability

The data underlying this article will be available in NCBI under Bioproject *tbd* and Zenodo at http://doi.org/10.5281/zenodo.4517398, in addition to that available in the Supplementary Material, upon publication. Raw sequence data will be deposited in the NCBI Short Read Archive (SRA accession *tbd*). Analysis scripts are available on Github at https://github.com/ryankschott/.

## Supplementary Figures and Tables

**Supplementary Figure 1.**
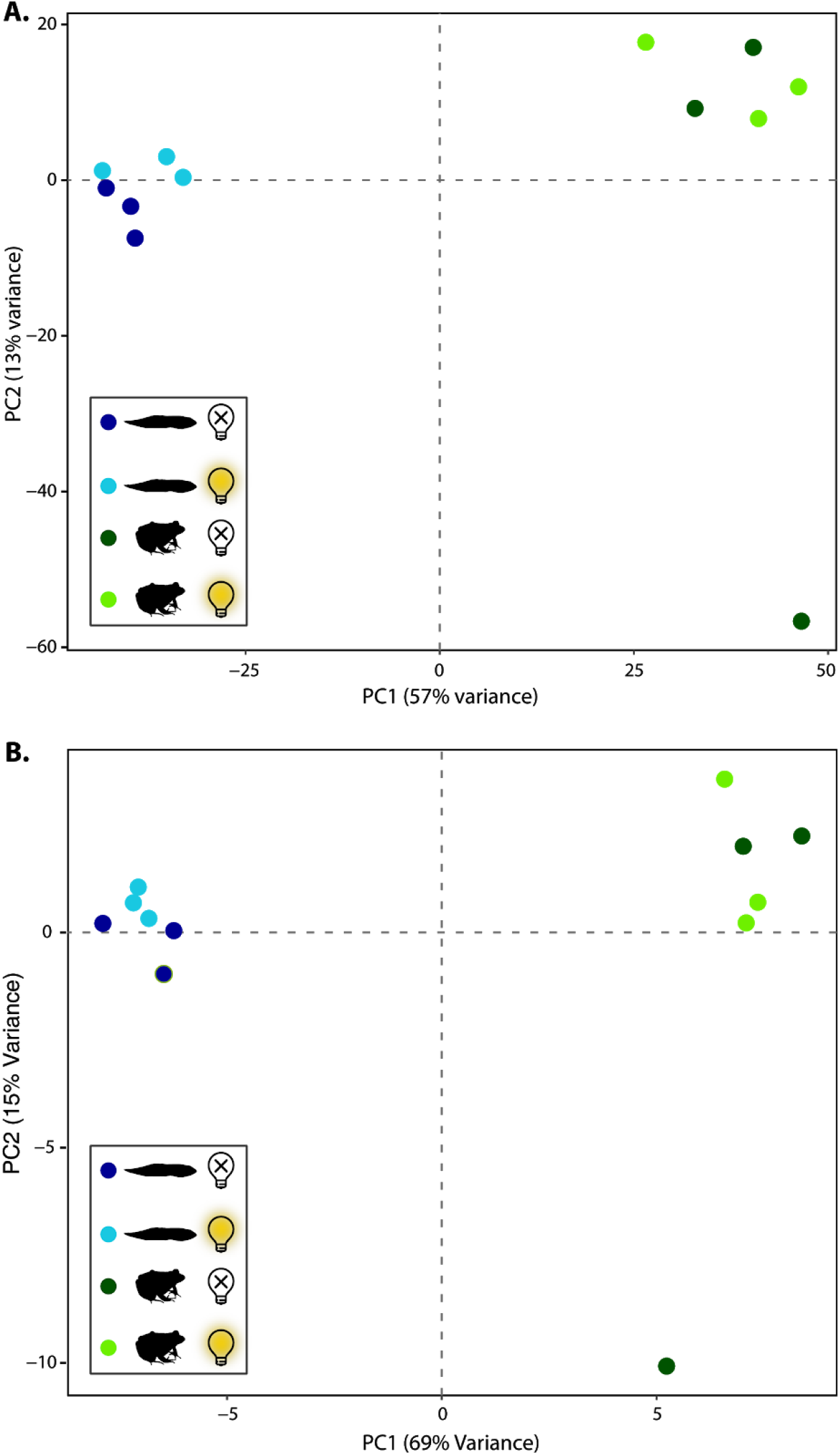
Principal components analysis plot of rlog transformed counts from the **A**) transcriptome-wide and **B**) vision gene coding sequence analyses illustrating the outlier juvenile, dark exposed sample. This sample was removed from subsequent analyses.

**Supplementary Figure 2.**
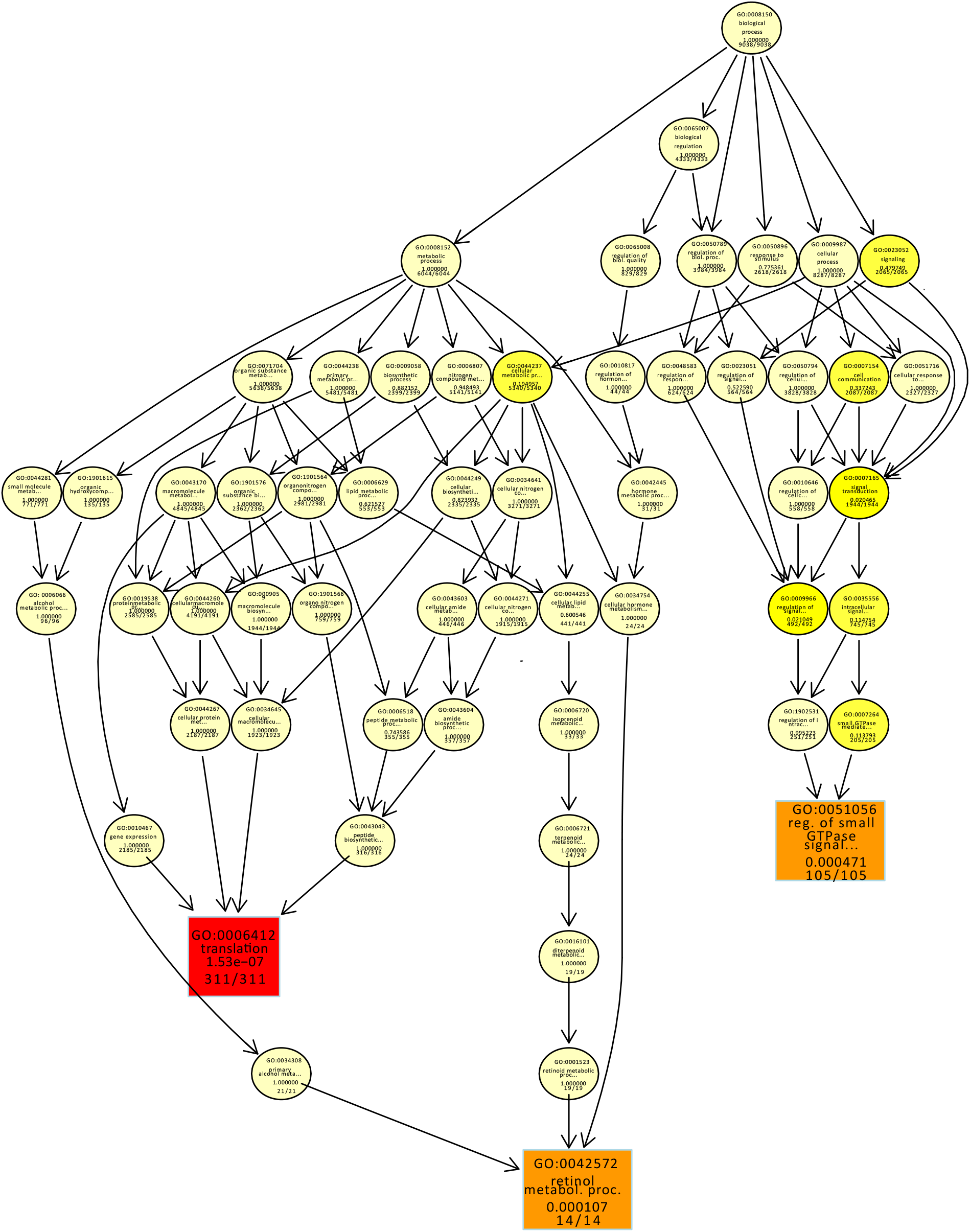
Enrichment of Gene ontology (GO) terms. GO terms, were based on annotation to the reduced set of transcripts using DIAMOND against the *Xenopus tropicalis* ENSEMBL database. GO enrichment analyses were performed using TopGO for the biological process gene ontology with the combined gene elimination and weighting algorithm (weight01) and Kolmogorov-Smirnov testing (KS).

**Supplementary Figure 3.**
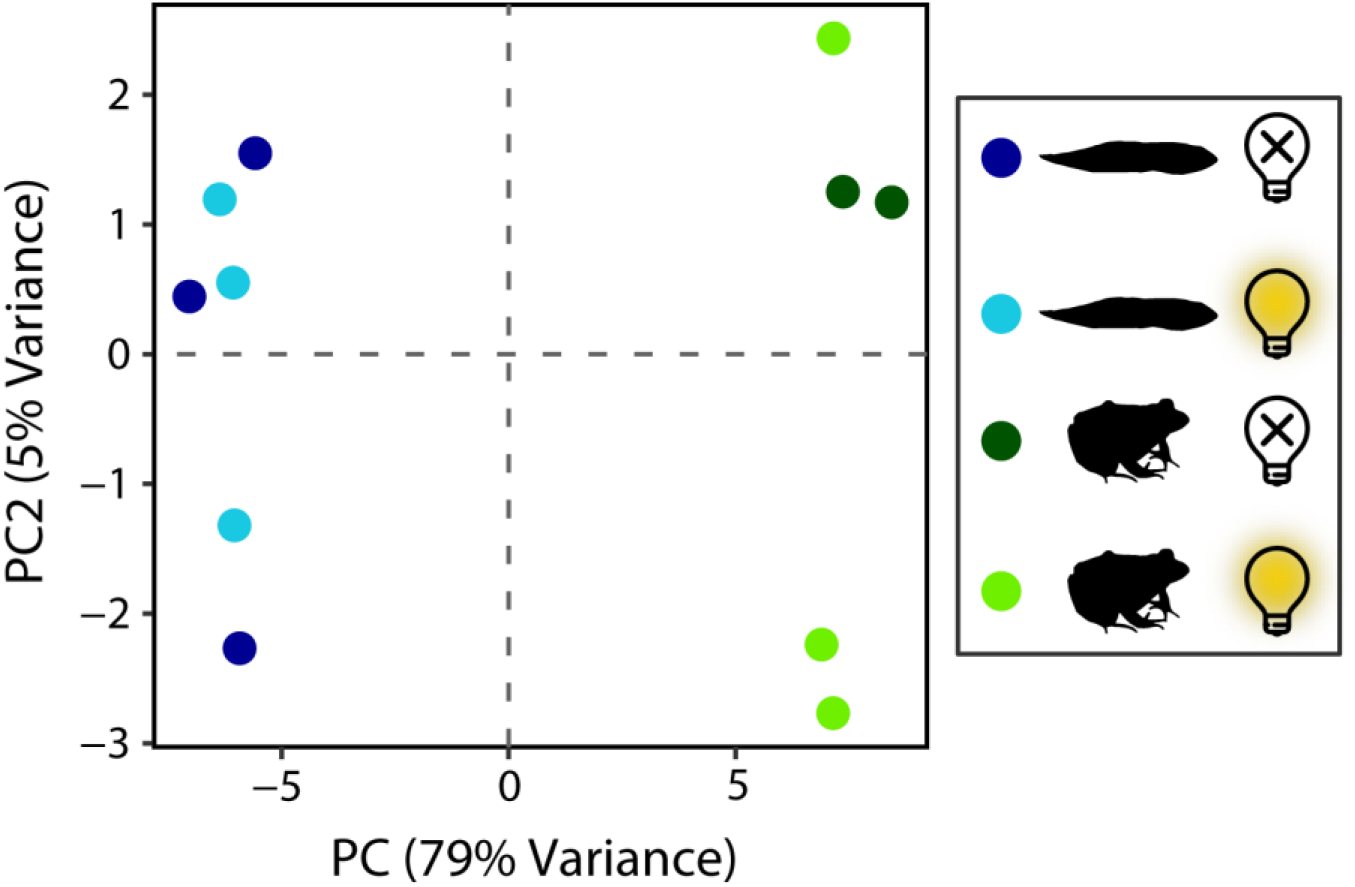
Principal components analysis plot of rlog transformed counts of visual gene coding sequences. The first principal component (PC1) accounting for 79% of the variance clearly separates juveniles and tadpoles. Light and dark exposure are not clearly separated by PC2, which accounts for only 5% of the variance.

**Supplementary Figure 4.**
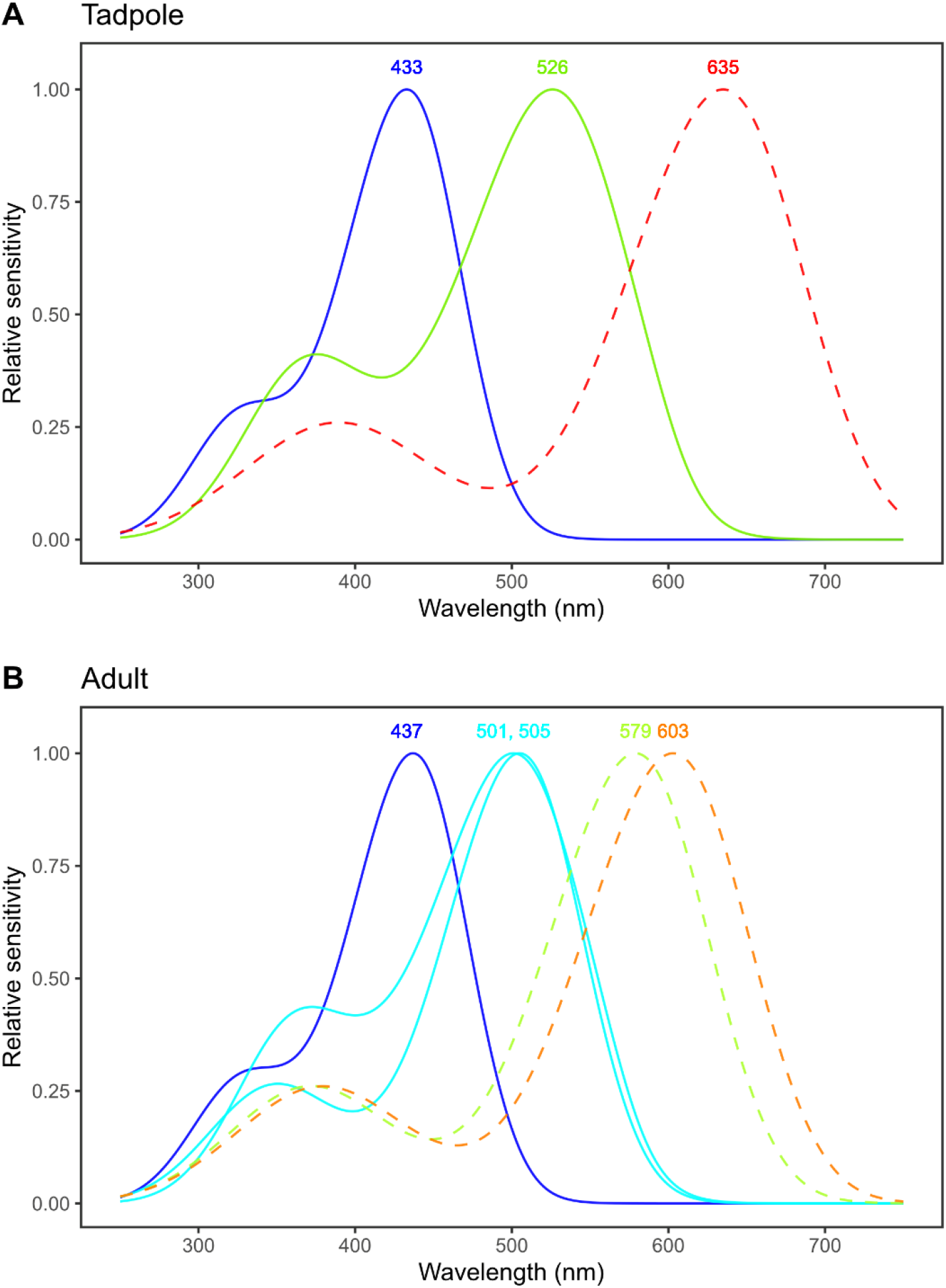
Spectral sensitivity of photoreceptors detected through microspectrophotometry in (**A**) one tadpole and (**B**) one adult *Lithobates sphenocephalus*. Govardovskii (2000) visual pigment templates for the mean λ_max_ and best-fit chromophore type found for each pigment are displayed. Data from rods are shown with solid lines, and cones with dashed.

**Supplementary Figure 5.**
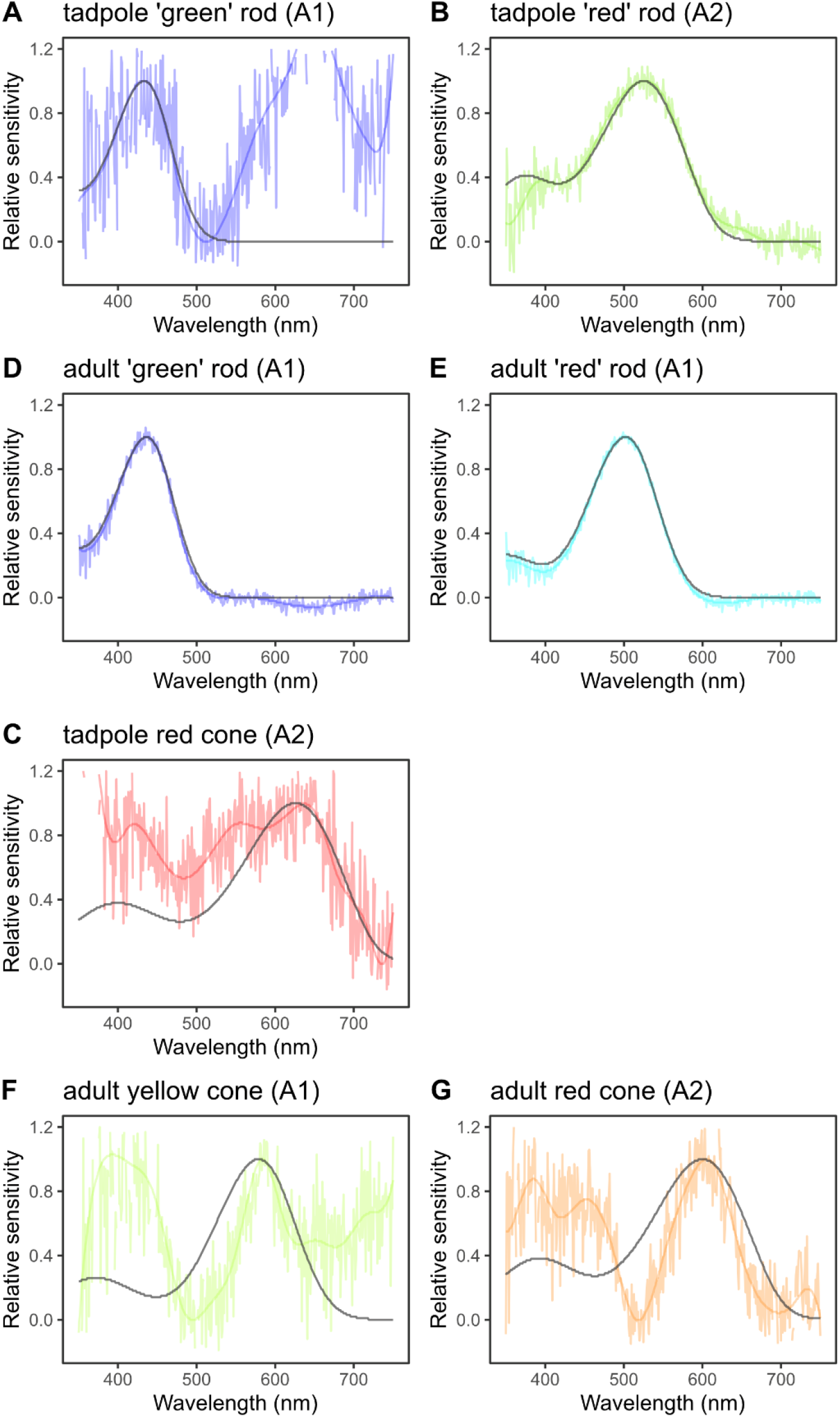
Examples of raw microspectrophotometry data and raw Gaussian fits (coloured and transparent) and Govardovskii (2000) template fits (black and non-transparent) for each cell and pigment type found in the tadpole (**A–C**) and adult (**D–G**) *Lithobates sphenocephalus*.

**Supplementary Figure 6.**
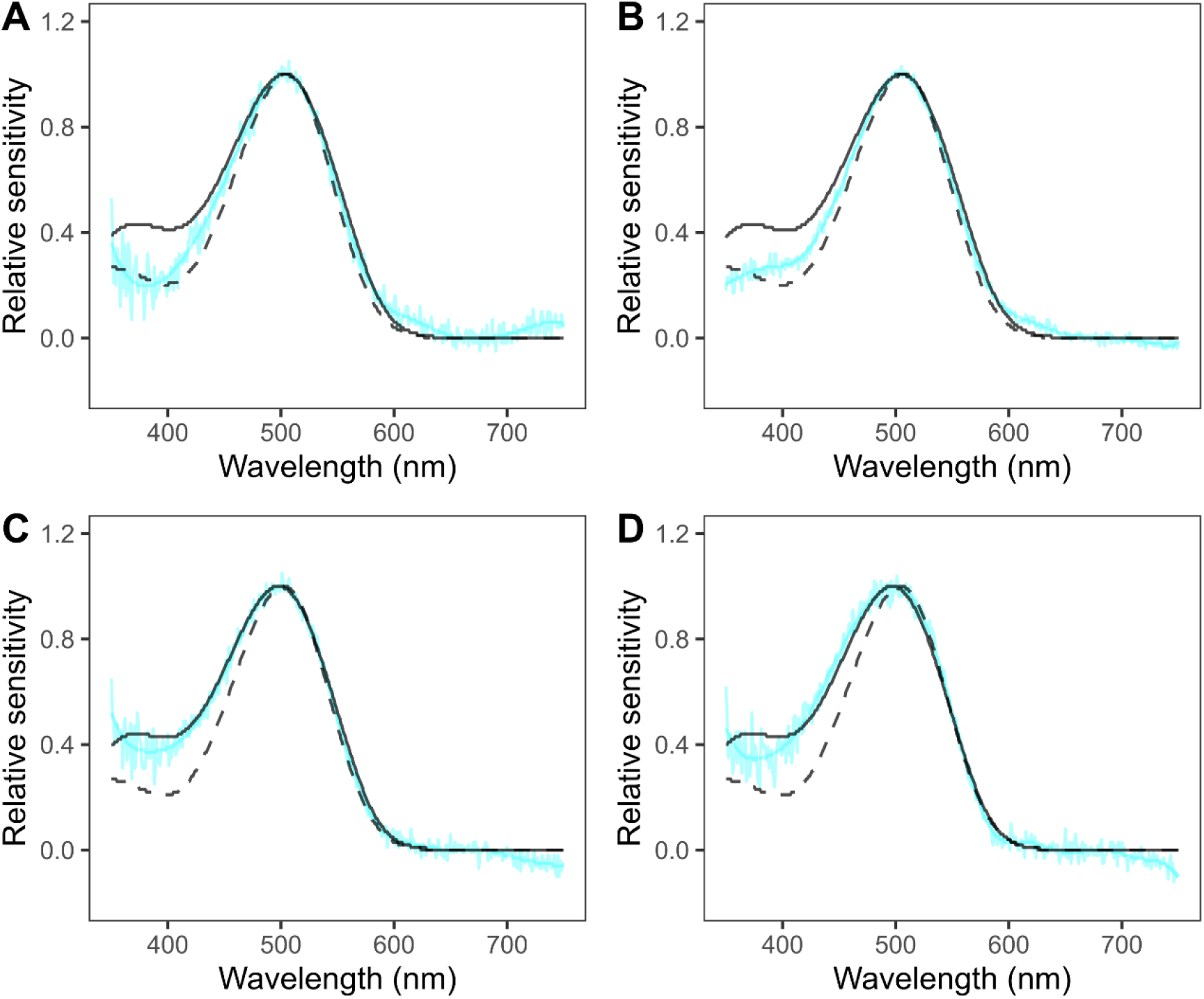
Examples of raw microspectrophotometry data and raw Gaussian fits (cyan) and Govardovskii (2000) template fits (black) for RH1 ‘red’ rods in *Lithobates sphenocephalus* adults where the A2 template (solid) was a better fit than the A1 template (dashed).

**Supplementary Figure 7.**
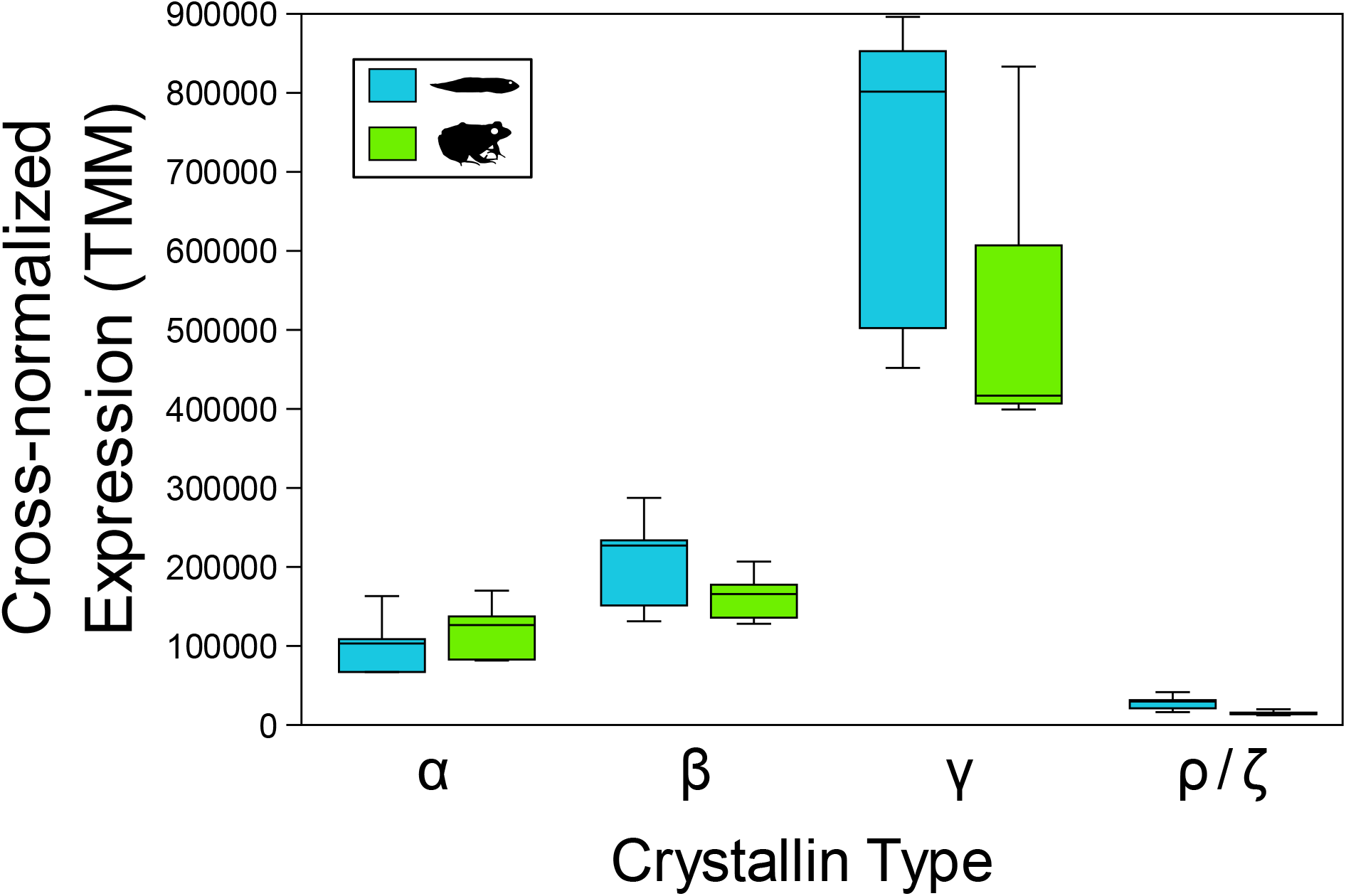
Comparison of averaged cross-normalized expression levels (TMM) for each major crystallin type. See also Supplementary Table 3 and Supplementary File 6.

**Supplementary Figure 8.**
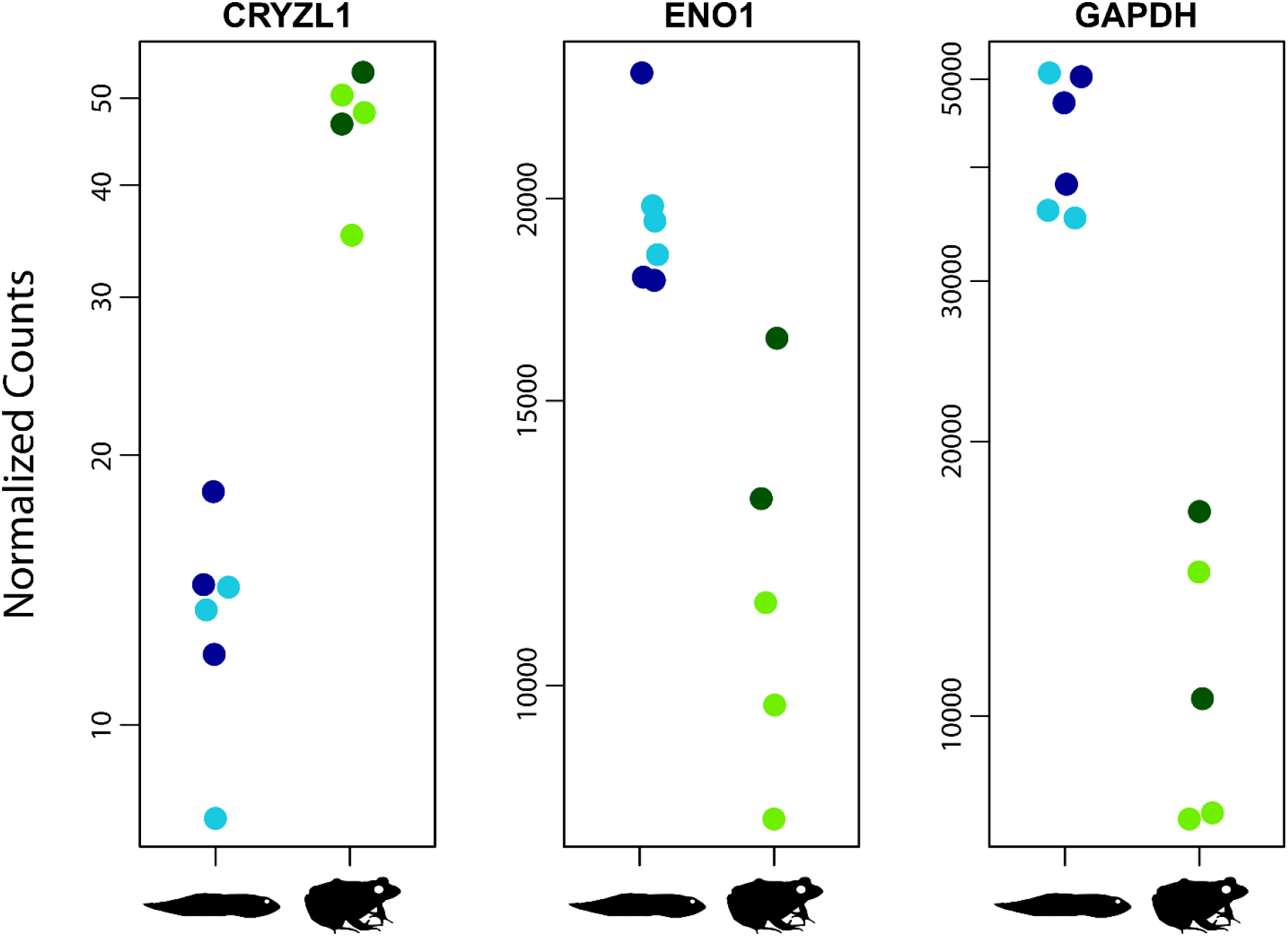
Expression profiles of taxon-specific lens crystallin genes that have not specifically been identified in frogs and that differ substantially between tadpole and adults (adjusted p-value < 0.05; Supplementary File 4). Plots are of normalized read counts for each gene.

**Supplementary Figure 9.**
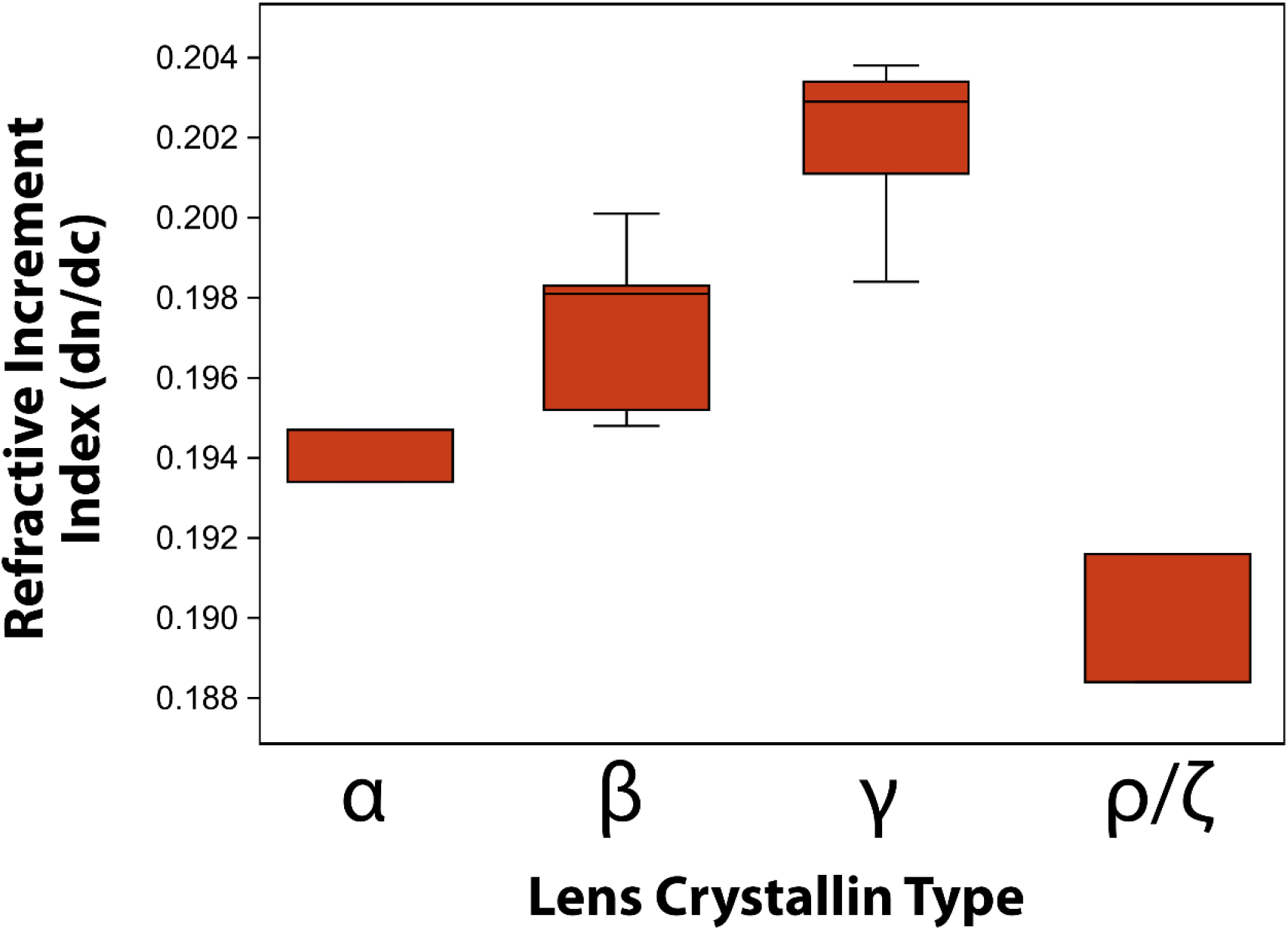
Comparison of averaged refractive increment index (*dn*/*dc*) for each major crystallin type. Values for *dn*/*dc* were computationally estimated using the Sedfit program (see Methods, Supplementary Table 3, and Supplementary File 6).

**Supplementary Table 1.** Sample information and read counts for the frogs used in the RNA-seq analyses. Sample MKF956 was identified as an outlier and was dropped from the analyses.

| Catalog No | Field No | Stage | Size (SVL)<br>or Gosner<br>Stage | Treatment | Raw Read<br>Count | Cleaned<br>Read<br>Count |
| --- | --- | --- | --- | --- | --- | --- |
| UTA *tbd* | MKF939 | Tadpole | 25–26 | Light | 23477325 | 14685429 |
| UTA *tbd* | MKF940 | Tadpole | 30–34 | Light | 28579708 | 17590719 |
| UTA *tbd* | MKF941 | Tadpole | 30–32 | Light | 30730778 | 20181240 |
| UTA *tbd* | MKF943 | Juvenile | 34.59 mm | Dark | 33986453 | 23719036 |
| UTA *tbd* | MKF944 | Juvenile | 25.64 mm | Light | 30326831 | 20244107 |
| UTA *tbd* | MKF945 | Juvenile | 24.77 mm | Light | 28315466 | 19763406 |
| UTA *tbd* | MKF946 | Juvenile | 26.30 mm | Light | 31367780 | 22952832 |
| UTA *tbd* | MKF948 | Tadpole | 25–27 | Dark | 32122782 | 21163991 |
| UTA *tbd* | MKF949 | Tadpole | 35–38 | Dark | 40393269 | 25858753 |
| UTA *tbd* | MKF950 | Tadpole | 35–38 | Dark | 32209800 | 23040731 |
| UTA *tbd* | MKF955 | Juvenile | 29.70 mm | Dark | 32457207 | 22374608 |
| UTA *tbd* | MKF956 | Juvenile | 32.74 mm | Dark | 51226316 | 32448910 |

**Abbreviations—SVL**, snout vent length.

**Supplementary Table 2.** Summary data from fits of individual photoreceptor scans from one *Lithobates sphenocephalus* tadpole (n = 41) and one adult (n = 33) to visual pigment templates from Govardovskii et al. (2000). Data were fit to templates based on 100% vitamin-A1 and 100% vitamin-A2 chromophores, and the best fit is indicated, though λ_max_ and standard deviation of λ_max_ are reported for both template fits. Fits were averaged by spectral class (region of the visible spectrum where λ_max_ occurs) and cell type (rod or cone, identified by microscopy during MSP). The adult frog from which MSP data were collected is USNM 591950. Complete MSP results are in Supplementary File 5.

| Age | Spectral Class | Cell Type | Visual Pigment | Best Fit | A1 $\lambda_{\max}$ | SD<sub>A1</sub> | A2 $\lambda_{\max}$ | SD<sub>A2</sub> | # Scans |
| --- | --- | --- | --- | --- | --- | --- | --- | --- | --- |
| tadpole | blue | rod | SWS2 | A1 | 433 | -- | 431 | -- | 1 |
|  | green | rod | RH1 | A2 | 533 | 5.1 | 526 | 4.5 | 38 |
|  | red | cone | LWS | A1 | 635 | 3.5 | 626 | 0 | 2 |
| juvenile | blue | rod | SWS2 | A1 | 437 | 1.5 | 442 | 2.6 | 8 |
|  | green | rod | RH1 | A1 | 505 | 2.1 | 504 | 2.1 | 18 |
|  | green | rod | RH1 | A2 | 505 | 2.1 | 501 | 3.9 | 4 |
|  | red | cone | LWS | A1 | 603 | 0 | 598 | 2.8 | 2 |
|  | yellow | cone | LWS | A1 | 579 | -- | 579 | -- | 1 |

**Supplementary Table 3.**
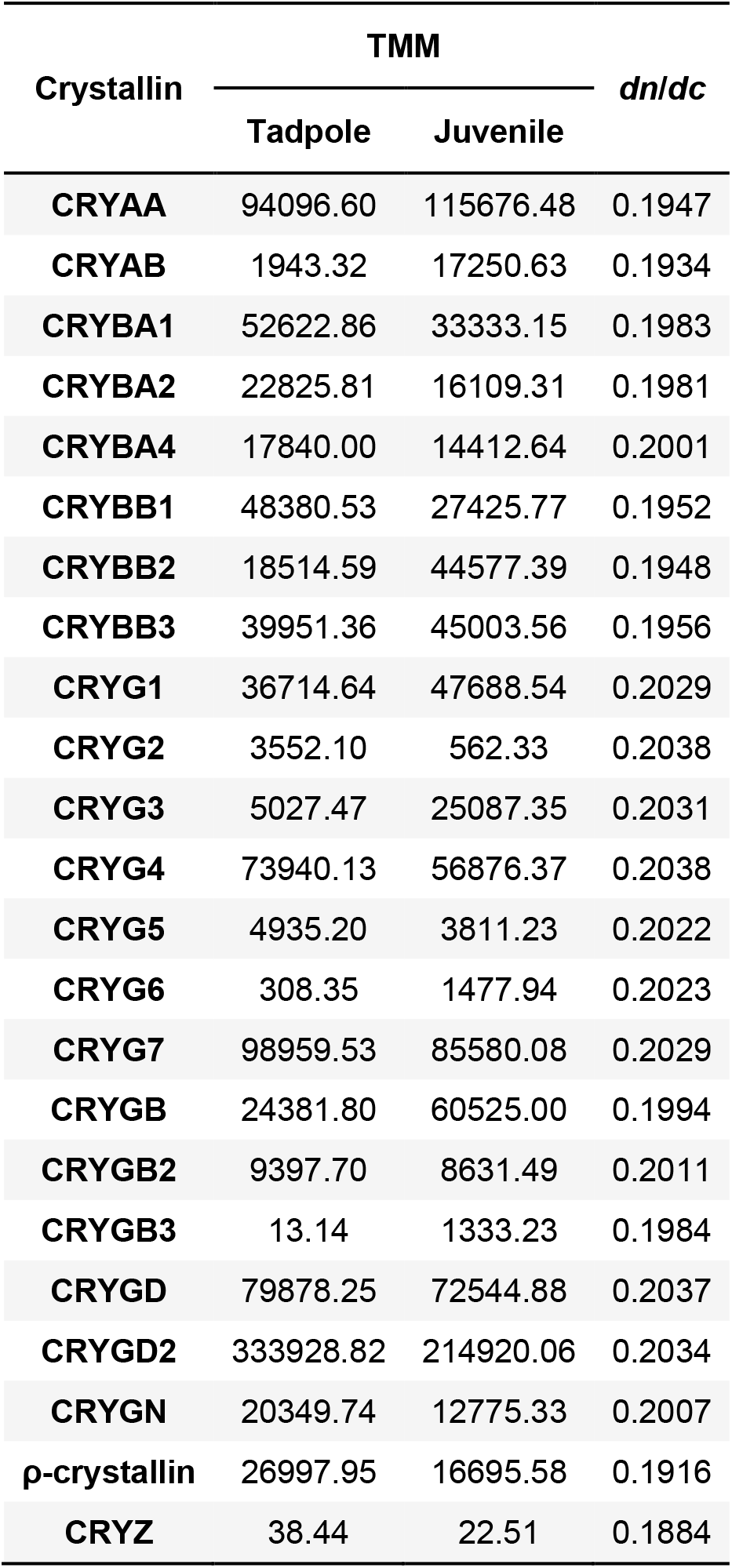
Cross normalized expression values (TMM) and computationally estimated refractive index increment (*dn*/*dc*) values for the ubiquitous and known frog taxon-specific specific crystallins. TMM values are averaged for the tadpole and juvenile replicates. A complete list of values can be found in Supplementary File 6.

